# Horizontal ‘gene drives’ harness indigenous bacteria for bioremediation

**DOI:** 10.1101/735886

**Authors:** Katherine E. French, Zhongrui Zhou, Norman Terry

## Abstract

Engineering bacteria to clean-up oil spills is rapidly advancing but faces regulatory hurdles and environmental concerns. Here, we develop a new technology to harness indigenous soil microbial communities for bioremediation by flooding local populations with catabolic genes for petroleum hydrocarbon degradation. Overexpressing three enzymes (almA, xylE, p450cam) in *E.coli* led to degradation rates of 60-99% of target hydrocarbon substrates. Mating experiments, fluorescence microscopy and TEM revealed indigenous bacteria could obtain these vectors from *E.coli* through several mechanisms of horizontal gene transfer (HGT), including conjugation and cytoplasmic exchange through nanotubes. Inoculating petroleum-polluted sediments with *E.coli* carrying the vector pSF-OXB15-p450camfusion showed that the *E.coli* die after five days but a variety of bacteria received and carried the vector for over 60 days after inoculation. Within 60 days, the total petroleum hydrocarbon content of the polluted soil was reduced by 46%. Pilot experiments show that vectors only persist in indigenous populations when “useful,” disappearing when this carbon source is removed. This approach to remediation could prime indigenous bacteria for degrading pollutants while providing minimal ecosystem disturbance.

## Introduction

Oil spills in recent decades have left a long-term mark on the environment, ecosystem functioning, and human health.^1–3^ In the Niger Delta alone, the roughly 12,000 spills since the 1970s have left wells contaminated with benzene levels 1000x greater than the safe limit established by the World Health organization and have irreparably damaged native mangrove ecosystems.^4,5^ Continued economic reliance on crude oil and legislation supporting the oil industry mean that the threat of spills is unlikely to go away in the near future.^6^

At present, there are few solutions to cleaning up oil spills. Current approaches to removing crude oil from the environment include chemical oxidation, soil removal, soil capping, incineration, and oil skimming (in marine contexts).^7,8^ While potentially a ‘quick fix,’ none of these solutions are ideal. Soil removal can be costly and simply moves toxic waste from one site to another.^9^ Chemical oxidants can alter soil microbial community composition and pollute groundwater.^10^ Incineration can increase the level of pollutants and carbon dioxide in the air and adversely affect human health.^11^ Practices such as skimming only remove the surface fraction of the oil while the water-soluble portion cannot be recovered, negatively effecting marine ecosystems.^12,13^

Synthetic biology has now given us the tools to tackle grand environmental challenges like industrial pollution and could usher in a new era of ecological engineering based on the coupling of synthetic organisms with natural ecosystem processes.^14–18^ Consequently, using bacteria specially engineered to degrade petroleum could present a viable solution to cleaning up oil spills in the near future. Previous studies have identified which bacterial enzymes are involved in petroleum hydrocarbon degradation (reviewed in references ^9,19^) and have engineered bacterial enzymes like p450cam for optimal *in vivo* and *in vitro* degradation of single-substrate hydrocarbons under lab conditions. ^20,21^ However, there are several critical gaps in or knowledge of engineering bacteria for oil-spill bioremediation. First, we know little about how the performance of these enzymes compare and which enzyme would present an ideal target for over-expression in engineered organisms. Second, it is unclear how well engineered organisms can degrade petroleum hydrocarbons compared to native wild-type bacteria which naturally degrade alkanes, such as *Pseudomonas putida*. Third, the environmental effects of engineered bacteria on native soil populations are unclear. For example, do these bacteria persist over time in contaminated soils? Although the use of genetically modified bacteria in bioremediation is attractive, this solution faces significant regulatory hurdles which prohibit the release of genetically modified organisms in the environment.^22^

Here, we propose a new bioremediation strategy which combines synthetic biology and microbial ecology and harnesses natural processes of horizontal gene transfer in soil ecosystems. We screened five enzymes involved in petroleum degradation in *E. coli* DH5α (alkB, almA, xylE, ndo and p450cam) to identify 1) where these enzymes localize and their effect on crude oil using advanced microscopy and 2) to asses each enzyme’s ability to degrade three petroleum hydrocarbon substrates (crude oil, dodecane, and benzo(a)pyrene) compared to two wild type bacteria (*Pseudomonas putida* and *Cupriavidus* sp. OPK) using bioassays and SPME GC-MS. Based on these results, we selected one vector (pSF-OXB15-p450camfusion) to determine whether small, synthetic vectors carrying catabolic genes could be transferred to indigenous bacteria found in petroleum-polluted sediments and whether this shift in community metabolism could increase rates of pollutant degradation.

## Results and Discussion

### Overexpression of petroleum hydrocarbon-degrading enzymes in E. coli

To compare the localization and activity of known petroleum hydrocarbon-degrading enzymes, we inserted five enzymes (alkB, almA, xylE, ndo, and p450cam) and required electron donors into the vector backbone pSF-OXB15 using Gibson Assembly^23^ (**SI Fig. 1**). To identify where each enzyme localized within *E. coli* DH5α, we tagged each enzyme with a fluorophore (gfp or mcherry). Fluorescence microscopy revealed that alkB was localized to bacterial cell membranes and almA was found throughout the cytoplasm. The camphor-5-monooxygenase camC from the p450cam operon was expressed throughout the cell cytoplasm while another enzyme in the operon, the 5-exo-hydroxycamphor dehydrogenase camD, was expressed within a small compartment at one end of the cell (**Fig. 1A**). The dioxygenase ndoC from the ndo operon was also localized to a small compartment at one end of the cell. The dioxygenase xylE was found in small amounts in the bacterial cell membrane and larger amounts in a microcompartment at one end of the cell. In all cases, these compartments were 115-130 nm wide and could be seen in young, mature and dividing cells. The presence of microcompartments in *E. coli* expressing p450cam, ndo, and xylE could reflect the known use of protein-based microcompartments by bacteria to concentrate highly reactive metabolic processes.^24^

**Fig. 1.**
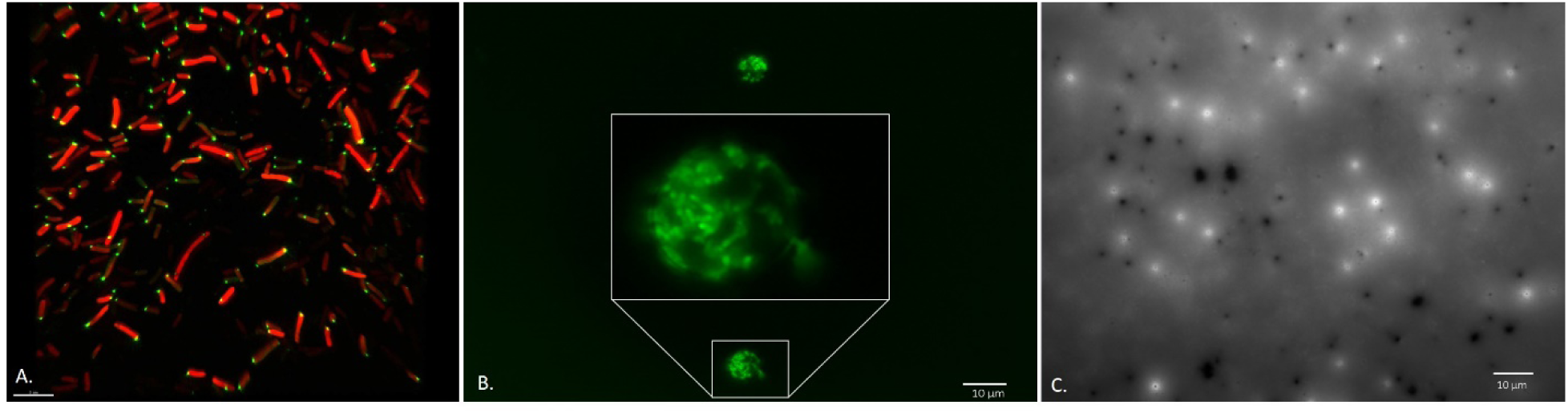
Expression and localization of bacterial monooxygenases and dioxygenases involved in petroleum degradation in *E*.*coli* DH5α. A. Structured Illumination Microscopy (SIM) image of *E.coli* DH5α expressing proteins involved in petroleum degradation (cam A,B, C and D) from the CAM plasmid in *E.coli*. camC (fused to mcherry) was found throughout the cell while camD (fused to gfp) localized to a microcompartment at one end of the cell. The scale bar is 5 μm. B. *E. coli* DH5α expressing alkB fuse to gfp were found clinging to spheres containing crude oil, mimicking a behavior seen in wild-type oil-degrading bacteria. C. EPS from *E. coli* DH5α expressing xylE. Gfp-tagged xylE were found around small pores (ca. 500 nm) within the EPS matrix. Figures 1B and 1C were taken using the GFP filter on a Zeiss AxioImager M1.

Over-expression of all five enzymes imbued *E. coli* DH5α with metabolism-dependent chemotactic behavior, where cell movement is driven towards substrates affecting cellular energy levels.^25^ *E.coli* DH5α do not have flagella, but rather, moved towards petroleum hydrocarbon substrates via twitching.^26–28^ Fluorescence microscopy showed *E. coli* DH5α expressing alkB ‘clinging’ to oil droplets (**Fig. 1B**) and those expressing xylE seemed to use the compartment-bound enzyme as a ‘guide’ towards crude oil (**SI Fig. 2**). Both behaviors mimic the interactions of wild-type oil-degrading bacteria.^7^

**Figure 2.**
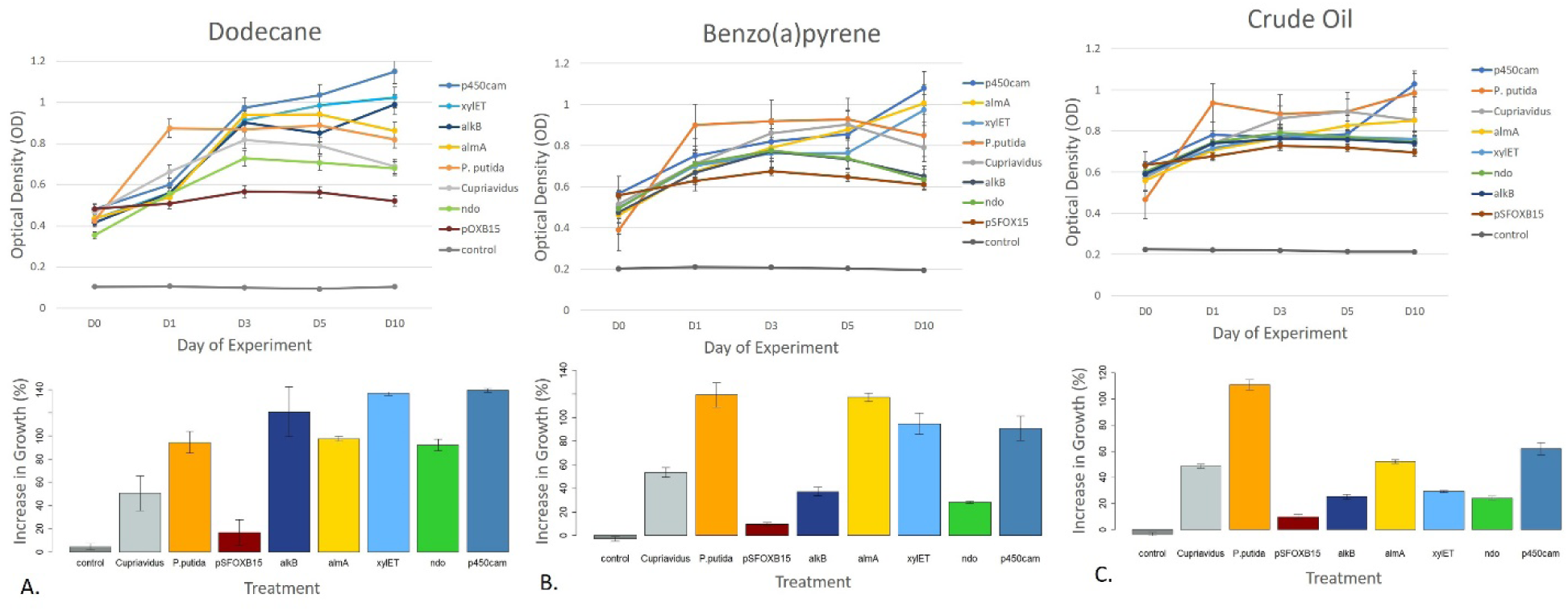
Growth of wild-type and engineered bacteria on dodecane (A), benzo(a)pyrene (B), and crude oil (C). Wild type strains are denoted as ‘*Cupriavidus*’ and ‘*P. putida*.’ Synthetic strains are denoted according to what enzyme they are engineered to express (e.g. alkB, almA). The two negative controls are a control with the carbon substrate but no cells and *E. coli* DH5α transformed with the vector backbone used in the experiment (pSFOXB15) but without genes inserted for hydrocarbon degradation.

Fluorescence microscopy also revealed for the first time the key role of extracellular enzymes in degradation of petroleum hydrocarbons. Three enzymes, alkB, almA, and p450cam were found in extracellular vesicles ranging in size from 0.68 μm to 1.67 μm (**SI Fig. 3**). These vesicles were only seen when *E. coli* DH5α was exposed to petroleum hydrocarbons. They are larger than minicells (which range from 200-400 nm in diameter)^29^ and seem to serve some other function. Confocal images suggest that these vesicles may come into contact with oil droplets, potentially attaching to (or merging with) their surface (**SI Fig 4**).

**Figure 3.**
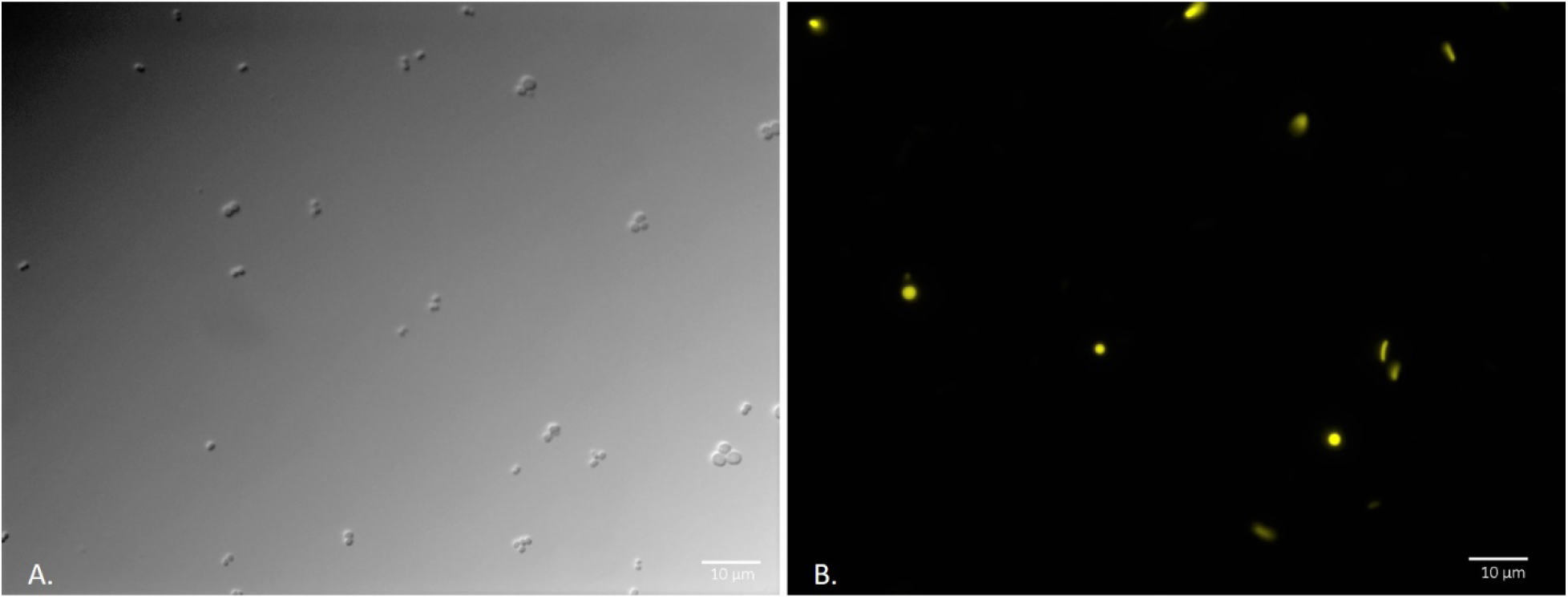
Expression of pSF-OXB15-p450camfusion in the marine bacteria *Planococcus citreus*. A. DIC image of wild-type *P. citreus* (no exposure to *E.coli*). These bacteria are coccoid-shaped with cells found individually or in groups of 1-4. B. Image of *P. citreus* and *E.coli* expressing p450cam after 48 hours of co-cultivation (described in methods). Fig. 3B was taken using the Texas Red filter on a Zeiss AxioImager M1 (excitation/emission 561/615).

**Figure 4.**
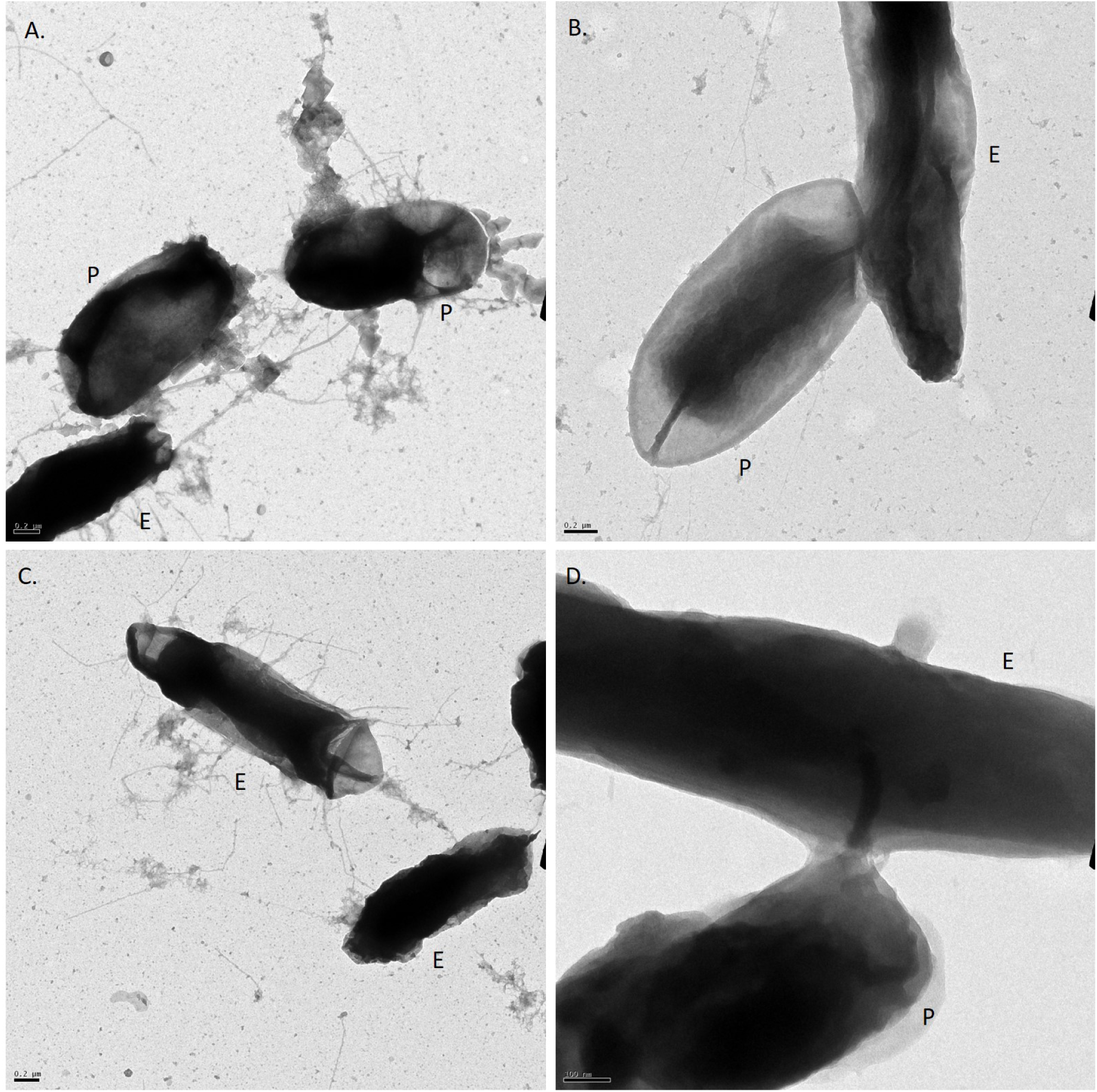
TEM confirmation of horizontal gene transfer among *E. coli* expressing pSF-OXB15-p450camfusion and *P. putida*. A. *E. coli* DH5α (E) connected to *P. putida* (P) via conjugative pili. B. Mating-pair bridge between *P. putida* and *E. coli* DH5α. C. *E. coli* DH5α with conjugative pili. D. Nanotube connecting *P. putida* and *E.coli* DH5α.

We also found three enzymes, alkB, xylE, and p450cam, within the *E. coli* exopolysaccharide (EPS) matrix. AlkB and xylE were concentrated around the 500 nm pores within the EPS and found dispersed in smaller amounts throughout the exopolysaccharide. In contrast, p450cam was distributed in high amounts throughout the EPS (**Fig. 1C**). Cryotome sectioning of the EPS from bacteria expressing p450cam indicates that the monooxygenase camA from the p450cam operon co-localized with a second enzyme involved in hydrocarbon degradation, the dehydrogenase camD (**SI Fig. 5**). Protein levels of EPS from bacteria expressing alkB (0.84 ± 0.04 mg/mL), xylE (0.85 ± 0.13 mg/mL), and p450cam (0.97 ± 0.06 mg/mL) were also significantly higher than *E. coli* DH5α expressing the empty vector pSF-OXB15 (0.44 ± 0.02 mg/mL) (**SI Fig 6**) and were comparable to the protein level of *Cupriavidus* sp. OPK (1.2 ± 0.15 mg/mL), a bacteria known to use biofilms to degrade crude oil.^7^ Although previous studies suggest that EPS may be involved in the extra-cellular metabolism of environmental pollutants,^7,30–34^ this is the first study to identify several enzymes which may play a role in this process.

**Figure 5.**
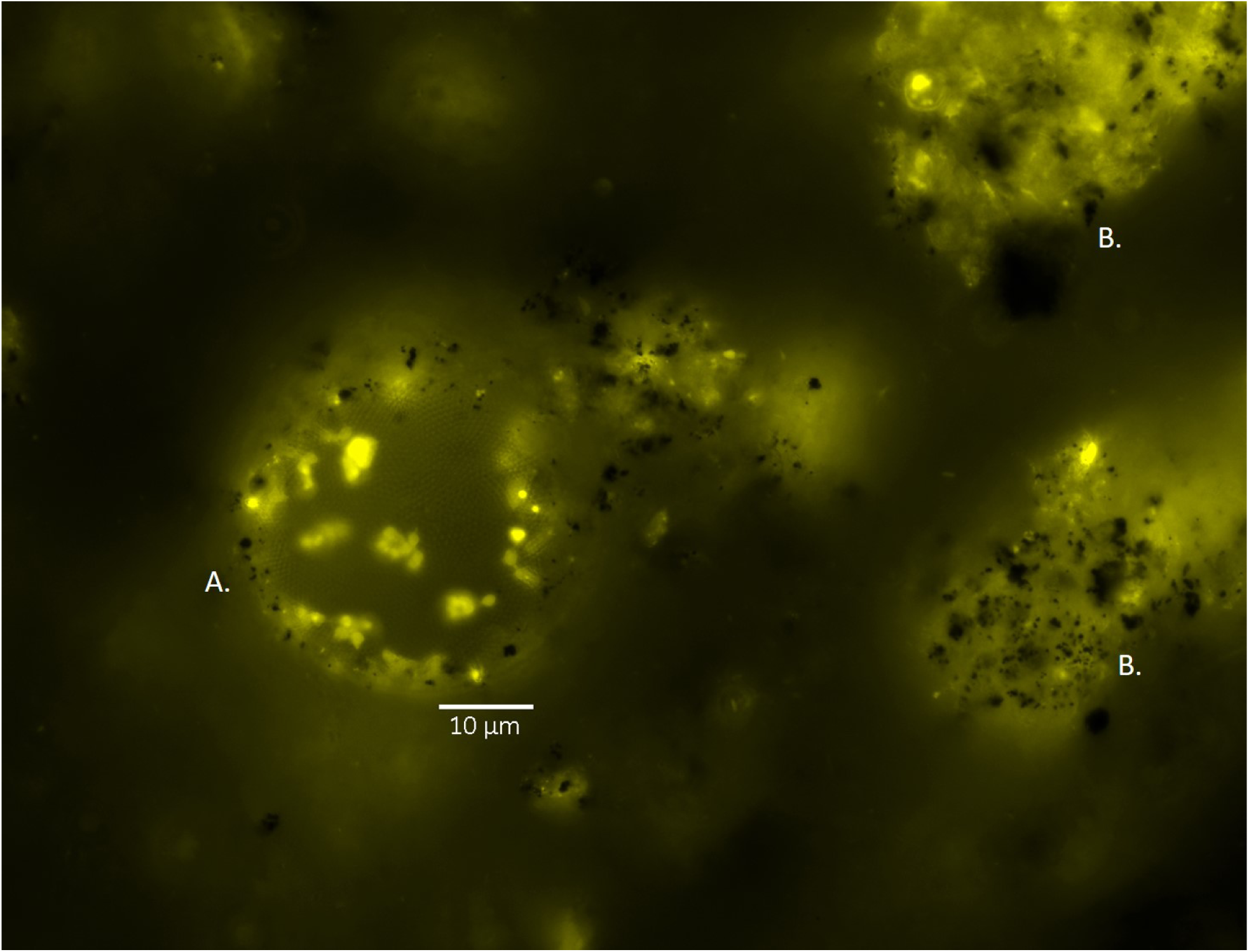
Expression of vector pSF-OXB15-p450cammcherry in indigenous microbial communities in sediment from a former Shell Oil refinery (Shell Pond, Bay Point, CA). A. Example of native soil bacteria expressing the vector. These bacteria construct spheres made of polymers in a honey-comb pattern. B. Expression of p450cam tagged with mcherry in biofilm matrices within sediment from Shell Pond. The biofilms form thin nets over large soil particles; small soil particles can be seen as black ‘dots’ sticking to the biofilm surface. Bacteria carrying the vector can be seen embedded in the EPS. In all microscopy images, mcherry is false-colored yellow. The image was taken using the Texas Red filter on a Zeiss AxioImager M1 (excitation/emission 561/615).

**Figure 6.**
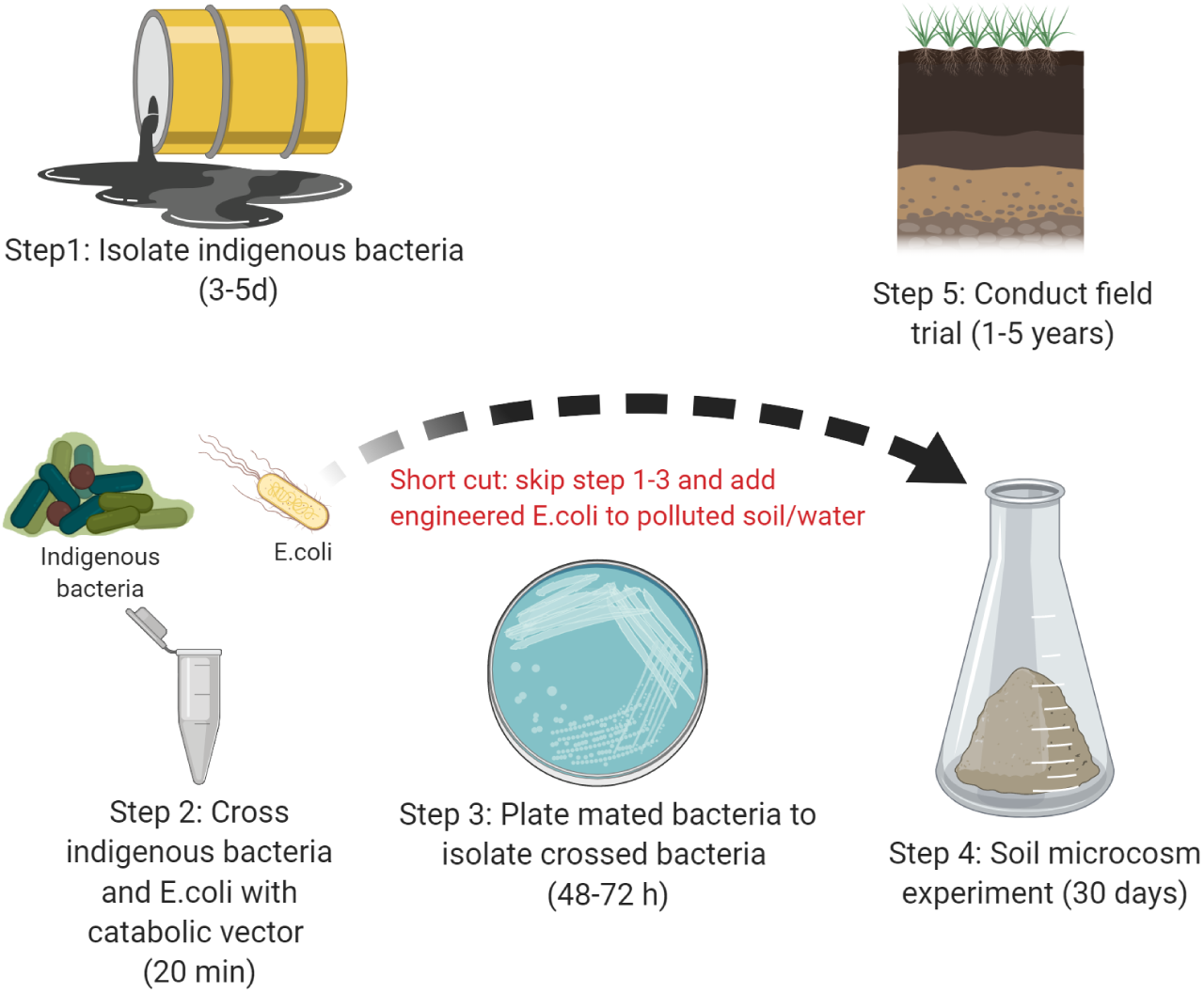
Overview of horizontal “gene-drive” system. The diagram depicts two approaches to flooding indigenous bacterial populations with catabolic genes of interest 1) through controlled mating and re-release or 2) direct application of *E.coli* DH5α with the catabolic genes of interest. Step 4 is there as a check to ensure the native bacteria take up the plasmid while in a soil matrix and can be shortened/extended as needed. The final step 5 is for monitoring purposes and can be shortened/extended as needed. This protocol works with water/marine samples (substituting water for soil).

Finally, extracellular enzyme expression also influenced the size of the oil droplets within the cell culture media. For example, *E. coli* DH5α expressing xylE produced very small oil droplets (primarily >1 μm in diameter) of crude oil while those expressing alkB and almA produced droplets ranging in size from 1 μm-120 μm in diameter (**SI Fig. 7**). Although xylE was not seen in vesicles, confocal suggests that crude oil droplets can flow through pores in biofilms and may become coated in EPS in the process. Potentially, attachment of enzymes to oil droplets (through fusion with vesicles or contact with EPS) may influence how fast a droplet is degraded over time. Previous studies have shown that vesicles embedded with enzymes can catalyze chemical reactions.^35,36^ In addition, Dmitriev et al. have shown that two bacteria, *P. putida* BS3701 and *Rhodococcus* sp. S67, use vesicular structures containing oxidative enzymes which attach to and play a role in degradation of crude oil droplets.^37^ Our results thus suggest that bacterial monooxygenases and dioxygenases involved in petroleum hydrocarbon degradation may be involved in multiple, complex inter and intra-cellular processes that lead to the degradation of crude oil.

### Comparison of enzyme activity

To determine which enzymes were most useful for degrading long-chain hydrocarbons, PAHs, and crude oil, we conducted 96-well plate assays exposing wild-type and genetically engineered bacteria to 1% of dodecane, benzo(a)pyrene or crude oil. We found that bacteria engineered to over-express specific enzymes in petroleum degradation were able to degrade single-carbon substrates better than the wild-type bacteria *P. putida* and *Cupriavidus* sp. OPK. ANOVA of the assay data showed that there was significant variation in bacterial growth when exposed to dodecane (F_8,31_ = 33.4, p = <0.001), benzo(a) pyrene (F_8,31_ = 73.03, p = <0.001) and crude oil (F_8,31_ = 240.6, p = <0.001). When exposed to dodecane, *E. coli* DH5α expressing p450cam increased in biomass the most (139.4%), followed by *E. coli* DH5α expressing xylE (136.3%), alkB (120.8%), and almA (97.6%) (**Fig. 2A**). Expressing p450cam and xylE led to significantly greater conversion of dodecane to biomass compared to *P. putida* (*t* = 4.71, df = 3.17, p < 0.01 and *t* = 4.41, df = 3.17, p < 0.01 respectively). SPME GC/MS analysis of these cultures revealed that all three bacteria degraded 99% of dodecane in 10 days. When exposed to benzo(a)pyrene, *P. putida* had the greatest increase in biomass (119.2%) followed by *E. coli* DH5α expressing almA (117.1%), xylE (94.8%), and p450cam (90.8%) (**Fig. 2B**). T-tests showed there was no significant difference in the biomass of *P. putida* and these three strains (p > 0.10). SPME GC-MS showed that *E.coli* expressing P450cam, almA and xylE degraded 90%, 97% and 98% of the benzo(a)pyrene respectively while *P. putida* degraded 86%.

In contrast, when engineered and wild-type bacteria were exposed to crude oil, *P. putida* converted the oil to biomass more efficiently, increasing in biomass by 110.9% (**Fig. 2C**). Only two genetically engineered bacteria, *E. coli* DH5α expressing p450cam and almA, had comparable increases in biomass to *Cupriavidus* sp. OPK (61.93%, 52%, and 48.7% respectively). The assay was repeated with crude oil stained with Nile Red and rates of degradation were determined according to French and Terry^7^. *P. putida* degraded 79% of crude oil while *E. coli* DH5α expressing p450cam and almA degraded 64% and 60% respectively. *E. coli* expressing alkB, xylE, and ndo only grew ∼25% and degraded 35-40% of crude oil. The high performance of p450cam when exposed to crude oil likely reflects the enzyme’s known substrate promiscuity^38,39^ which makes it a better catalyst for degrading crude oil, a complex substrate made of over 1,000 compounds.^40^

### Vector-exchange between E.coli DH5α and indigenous bacteria

To determine whether our engineered bacteria could transfer non-conjugative, synthetic vectors containing petroleum-degrading genes to indigenous soil and marine bacteria, we conducted a series of mating experiments. We found that wild-type bacteria readily received the vector pSF-OXB15-p450camfusion through horizontal gene transfer (HGT) (**Fig. 3**; **SI Fig. 8**). Rates for transformation ranged from 19 to 84% in 48 hours depending on the recipient species (**SI Table 1**) and were >90% after seven days of incubation for all tested species. Plasmid expression was stable for over three months in the absence of antibiotic pressure. Although our vectors carried a ColE1 origin of replication, this did not seem to present a barrier to HGT. This agrees with previous studies which suggest ColE1 plasmids can be found in wild bacteria^41^ and wild-type bacteria can receive ColE1 plasmids from *E.coli*.^42^

HGT can occur through transformation, transduction, conjugation, transposable elements, and the fusing of outer membrane vesicles (OMVs) from one species to another.^43,44^ To test whether wild-type bacteria could take up naked plasmids from cell culture, we adding 1 μl of purified plasmid (at a concentration of 1 ng/μl and 10 ng/μl) to LB cultures containing wild-type bacteria and by spreading diluted vectors onto agar plates. We saw no transformed cells. In addition, neither fluorescence microscopy or TEM showed OMV production or release by the *E.coli* DH5α strains created in this study. OMVs are 50-250 nm in diameter,^45^ much smaller than any of the vesicles produced by our strains.

To determine whether HGT of the synthetic vectors to wild-type bacteria was achieved through mating, we conducted TEM of wild-type bacteria after exposure to *E.coli* DH5α carrying pSF-OXB15-p450camfusion. TEM suggests several mechanisms of HGT through direct cell-to-cell contact may explain how vectors were transferred between transgenic *E. coli* DH5α and wild-type bacteria.^46^ In our study, we found *E.coli* DH5α and wild-type bacteria engaging in DNA transfer through conjugation and cell merging and in cytoplasmic transfer via nanotube networks. TEM showed *E. coli* DH5α expressing p450cam tethered to wild-type cells by conjugative pili over long distances (**Fig 4A**), the formation of mating pair bridges between wild-type cells and *E.coli* DH5α (**Fig 4B**), *E.coli* with conjugative pili (**Fig. 4C**), and E.coli and wild-type cells connected via nanotubes (**Fig. 4D**) (see also **SI Fig. 9**). Although the plasmids used in this study were non-conjugative, such plasmids can be mobilized and transmitted via conjugation in the presence of other conjugative plasmids or by merging with conjugative plasmids.^47–49^ Previous studies have also shown that plasmids (and chromosomal DNA) can be transferred through cell-contact dependent transfer without the use of conjugative pilii. This mechanism was first observed in 1968 in *Bacillus subtilis* and subsequently in other species (e.g. *Vibrio, Pseudomonas, Escherichia*) (reviewed in^49^). For example, Paul et al.^50^ found that lab strains of *E.coli* could transfer plasmid DNA to *Vibrio* through this process. Nonconjugal plasmids can also be transferred from between bacteria through nanotubes.^51^ Dubey and Ben-Yehuda^52^ show in their classic paper that gfp molecules, calcein, and plasmids could be transferred between *B. subtilis* cells. They also show that a non-integrative vector carrying a resistance marker from *B. subtilis* could be transferred to *Staphylococcus aureus* and *E.coli* (with recipient cells expressing antibiotic resistance). This transfer was rapid and could happen in as little as 30 minutes. In our study, it is impossible to say definitively by which mechanism our vectors were transferred from *E.coli* DH5α to the wild-type bacteria and in reality multiple mechanisms of transfer may be possible.

We conducted an additional experiment to determine the survival rate of engineered bacteria in petroleum polluted sediment from a former Shell Pond refinery in Bay Point, CA and whether these genes could be transferred to native, complex soil microbial communities. This sediment is contaminated with high levels of petroleum hydrocarbons, arsenic, heavy metals, and carbon black. At D_0_, *E.coli* DH5α containing the plasmid pSF-OXB15-p450camfusion were seen in aliquots of contaminated sediment and there were no autofluorescent bacteria visible in the media. After D_5_, the population of *E.coli* DH5α had declined. Instead, a number of diverse native soil bacteria now contained the plasmid, a trend which continued over the course of the experiment (**Fig. 5**). Based on morphological analysis of over 100 microscopy images and pairwise mating between bacteria isolated and identified (via sequencing) from Shell Pond, these bacteria belonged to the *Pseudomonas, Flavobacteria*, and *Actinomycete* genera among others. Plating out aliquots of soil at regular time points confirmed the data gathered by microscopy: the number and diversity of bacteria expressing the plasmid increased 50-fold over the first 30 days of the experiment (from 2.6 ×10^−4^ CFU at D_0_ to 125 ×10^−4^ CFU at D_30_) (**SI Fig. 10**). The spider-silk-like biofilms formed by native soil microbiota present in the soil were also fluorescent (**Fig. 5**), suggesting the p450cam enzymes also play a role in extracellular degradation of petroleum hydrocarbons under real-world conditions. Preliminary GC/MS analysis showed that the amount of total petroleum hydrocarbons within the contaminated sediments decreased by 46% within 60 days. We left the experiment running and after 120 days bacteria carrying the vector were still prolific (**SI Fig. 11**). A pilot experiment using artificially contaminated water samples suggest that genes are transferred from *E.coli* to indigenous bacteria only when oil is present. In the control samples without oil, we saw no bacteria carrying the vector over the course of the 30-day experiment (the experiment will be published fully in a later publication).

HGT is thought to play a role in the degradation of environmental toxins^53^ and several studies have shown that wild-type bacteria carrying large plasmids with degradative genes can pass these genes on to a limited number of bacteria.^54^ However, this is the first study to provide evidence for HGT between *E.coli* DH5α carrying a small, non-conjugative vector and wild soil microbiota. Our results show that adding engineered *E.coli* DH5α carrying small synthetic plasmids to polluted environmental samples may be even more effective than adding a wild-type bacteria with a larger catabolic vector. Previous studies show that rates of HGT between the donor and recipient bacteria in soil are low (e.g. 3 x 10^−3^ CFU), recipient cells come from only a few genera, and the spread of the catabolic vector through the microbial community does not always lead to enhanced degradation.^55–57^ The high transfer rate of synthetic plasmids like the ones used in this study could be due to several reasons, including the small size of the vector (natural plasmids carrying degradative genes, e.g. OCT from *Pseudomonas putida*, are >30k bp), the inability of *E.coli* DH5α to avoid conjugation/cytoplasmic exchange with wild-type bacteria, and selective pressure (acquiring genes related to hydrocarbon degradation would increase host fitness).^58,59^ Our results also suggest native soil and marine microbial communities will continue to carry synthetic vectors with useful catabolic genes, likely until the metabolic cost of replicating the vector no longer presents an advantage. This agrees with previous studies which show that HGT transfer events that alter organism metabolism can increase organism fitness by allowing them to colonize new ecological niches.^57,60,61^ It also suggests that this approach to engineering polluted ecosystems is self-limiting: selective pressure from the existing ecosystem could remove introduced genes from local microbial populations when no longer needed.

Our results suggest transgenic bacteria can easily transfer genes to wild-type soil bacteria. Potentially, engineered bacteria could be used in soil and marine-based ‘gene drives.’ This drive could be achieved two ways: by adding the engineered *E.coli* DH5α carrying catabolic genes of interest directly to oil-contaminated soil/water *or* by adding pre-mated indigenous bacteria to the contaminated site (**Fig. 6**). Previous studies have proposed this form of ecological engineering,^62,63^ but the best of our knowledge no studies have shown that such drives would be successful. *E.coli* DH5α engineered to carry plasmids containing genes involved in degradation of environmental toxins could be used to augment the capacity of native soil microbial communities to degrade pollutants of interest. Replacing antibiotic selection markers with chromoprotein ones^64^ would eliminate the release of antibiotic resistance genes into the environment.

The primary barriers to implementing this approach on current polluted industrial sites are 1) lack of standardized procedures to test and ultimately allow the use of GM organisms for environmental applications and 2) the willingness of site managers to adopt this approach to remediation. The first barrier can be overcome by developing a comprehensive checklist system for assessing the impact of GM organisms on the environment which would involve a standardized field-trial with appropriate biocontainment measures in place. For example, soil cores could be taken, encased in plastic tubing, and returned to the site for long-term monitoring on the effects of the GM organism on microbial community structure, pollutant degradation, and other factors such as vegetation cover. The EPA does have a system in place for assessing the risk posed by GM bacteria to human and environmental health^65^; yet the assessment procedures used aren’t clear from publicly available documentation and the frequency of researchers seeking out EPA approval for field trial of bacteria engineered for remediation is unknown. Publicly available documentation and potentially even de-centralized approval of GM field trials (e.g. through university Environmental Health & Safety offices) could make field trials of GM bacteria more achievable in the near future. The second barrier can be overcome through public engagement with those working in the remediation sector (industry, site managers, and remediation consulting firms) and a shift in our approach to how we conduct remediation (favoring slower biological-based solutions that harness local ecological and chemical processes over faster processes such as oxidation and soil removal).

## Conclusion

Cleaning up environmental contamination from human activities is one of the greatest un-met challenges of the 21^st^ century. Our pilot research has shown that transferring catabolic genes involved in petroleum degradation from *E. coli* DH5α to indigenous bacteria may be a viable solution. This system could be adapted to exploit genes from local microbial populations which are already primed for degradation. This could be achieved by isolation and identification of native strains which degrade petroleum and proteomic identification and screening of candidate enzymes for over-expression. Future research is needed to determine 1) how long these plasmids are maintained under field conditions, 2) whether genetic mutations accumulate over time that might impact enzyme functioning, and 3) how vector-based gene drives harnessing natural processes of conjugation may affect local microbial community composition and soil metabolic functions. Concentrated efforts among microbiologists, ecologists, synthetic biologists and policy makers in this new area of research may usher in a new era of how we respond to environmental disasters and toxic waste management in the Anthropocene.

## Methods

### Vector design and cloning

All vectors were constructed using the vector backbone pSF-OXB15 (Oxford Genetics) and the Gibson Assembly cloning method. Gene sequences for all enzymes were gathered from UniProt (SI Table 2). Gene sequences were sent to Twist Biosciences for synthesis. All primers were synthesized by Genewiz (SI Table 3). We used the Q5 High-Fidelity Polymerase 2x Master Mix (New England Biolabs) to amplify genes and vectors under the following general conditions: 3 minutes at 95°C for denaturation, 30 cycles of 20s denaturation at 95°C, 30 seconds annealing at 60°C, and 30 seconds per kb for extension at 72°C, followed by one cycle of final extension for 3 minutes at 72°C. Annealing temperatures ranged from 50-65°C depending upon the gene amplified. Genes were assembled into the vector backbone using the New England Biolabs HiFi Assembly Master Mix. To transform *E.coli*, we added 5 μl of ligation reaction to DH5α cells (NEB), incubated cells on ice for 30 minutes, heat-shocked cells for 60 seconds, and allowed cells to recover for five minutes on ice. We added 950 μl of SOC media to each Eppendorf and placed cultures into a shaking incubator set to 30°C for one hour of vigorous shaking (250 rpm). We plated 100 μl of cell culture onto LB plates with kanamycin (50 μg/mL). Plates were incubated for 24-48 hours in an incubator set to 37°C. Colony PCR was performed following standard procedures to screen for colonies with the correct construct. Plasmids from positive constructs were verified using Sanger Sequencing through Genewiz and the UC Berkeley DNA Sequencing Facility.

### Verification of protein expression

Protein expression was confirmed by SDS-PAGE following standard procedures. Briefly, cell cultures containing the plasmids used in this study were grown overnight and 1mL aliquots were lysed using Cellytic tablets (Sigma-Aldrich). We mixed 30 μl of sample with 10 μl of 4x LDS loading buffer, boiled the mixture at 100° C for 2 minutes, and loaded 15 μl of each mixture into a Biorad MiniProtean TGX 12% precast gel. Gels were run on a Biorad MiniProtean Tetra system for 30 minutes, washed three times, immersed in 50 mL BioRad Biosafe Coomassie Dye for 1 hour, washed with water for 30 minutes, and then allowed to soak overnight in water to remove residual dye. Bands were observed the next day.

### Quantitative analysis of protein content in EPS

To quantify the total amount of protein found in the EPS of genetically engineered bacteria created in this study and control (pSF-OXB15) and wild-type (*P. putida, Cupriavidus* sp. OPK) strains, we conducted a Bradford assay following standard procedures for 96-well plate assays. Briefly, a calibration curve was made using BSA (Sigma-Aldrich) as a standard. Protein concentrations ranged from 0 to 1 mg/mL. Absorbance was detected ay 595 nm using a plate reader. This calibration curve was used to quantify the amount of crude protein in the EPS. Each treatment was replicated three times.

### Bioactivity assays

To determine the substrate specificity of each of the five enzymes (alkB, almA, xylE, ndo and p450cam) in vivo, we conducted 96-well plate assays using crude oil, dodecane and benzo(a)pyrene as model substrates. Briefly, each well contained 100 μl of minimal salt media, 100 μl of overnight cell culture, and 1% (2 μl) of hydrocarbon substrate. Each treatment was replicated four times. We used two positive controls (two known petroleum-degrading bacteria, *P. putida* and *Cupriavidus* sp. OPK) and two negative controls (*E. coli* with an empty vector backbone (pSF-OXB15) and no bacteria). OD readings were taken on a plate reader at 600 nm at four time points (D_0_, D_3_, D_5_, and D_10_). Each treatment was replicated four times.

### Nile Red Assay for determination of crude oil degradation

To determine the amount of crude oil degraded by bacteria, we used the Nile Red assay method described in French and Terry.^7^ Briefly, 0.1 mM stock solutions of Nile Red were made in DMSO. These stock solutions were encased in tin foil and kept in a −20°C freezer until use. To set up each assay, 96 well plates were filled with crude oil which had been complexed with Nile Red (10ul of Nile Red per mL of crude oil). For the control wells, MSM was added to the wells until a total of 200 μl was reached. For wells containing bacteria, 100 μl of bacteria in (OD of 0.6) were added to each well; MSM was then added to bring the total volume of each well to 200 μl. Each treatment was replicated three times. Assays were sealed with parafilm and placed on a shaking incubator (120 rpm) in a dark room. To measure FI and OD, we used a Tecan plate reader. We measured FI and OD at T_0_,T_1,_T_3_,T_5,_ and T_10_.

### GC/MS

Samples were removed from the 96-well plates and wells were washed with either 200 μl PBS (for dodecane) or 400 μl of acetone (for benzo(a)pyrene). Samples were placed into headspace vials (Agilent) and SPME headspace analysis was performed according to the method of Cam and Gagni^66^ with minor modifications. Samples were extracted for 30 minutes at 60°C. Analysis of total petroleum hydrocarbon content of contaminated sediment from Shell Pond was performed by Eurofins Calscience using EPA method 8015B (C6-C44).

### Mating experiments

To mate engineered (donor) and wild-type (recipient) bacteria, we mixed *E. coli* containing our plasmids with wild type bacteria (*P. putida, Cupriavidus* sp. OPK, *Planococcus citreus*, and *Microbacterium oxydans*) in a 1:1 ratio in 50 mL centrifuge tubes. Cultures were centrifuged at 2000 rpm for 10 minutes at room temperature. We removed the supernatant and resuspended the pellet in 40 μl of LB. We spotted this solution onto LB plates and allowed the colonies to grow for 48 h. One mixed colony was selected and resuspended in 1 mL of PBS. We plated 100 μl of the bacterial suspension onto LB plates with kanamycin and allowed them to grow overnight. To identify whether mating was successful, we selected single colonies and suspended them in 1 mL of SOC media in 2 mL Eppendorfs. Eppendorfs were then placed in a shaking incubator set to 250 rpm for 2 hours. Aliquots were removed from the Eppendorfs for microscopy. PCR amplification was used to determine the presence of the vector using Fwd 5’-TAACATGGCCATCATCAAGGAG-3’ and Rev 5’-ACCCTTGGTCACCTTCAG-3’. To determine whether wild-type bacteria would retain foreign plasmids with petroleum degrading genes in the absence of antibiotic selection pressure, we separated wild-type bacteria from *E. coli* by streaking for single colonies and maintained transformed wild-type bacteria in LB culture, changing the media twice a week for 12 weeks. To visually check for plasmid expression, 100 μl aliquots of culture were incubated on LB plates containing kanamycin every two weeks to check for colony growth.

### Soil survival experiment

To determine the survival rate of engineered bacteria in contaminated soils, we added *E.coli* containing the plasmid pSF-OXB15-p450camfusion to sediment taken from a former Shell refinery in Bay Point, CA. The sediment is composed of carbon black and contains petroleum hydrocarbons, heavy metals, arsenic, and other contaminants. We added 1 g of sediment to a 2 mL Eppendorf. We then centrifuged 500 μl of overnight bacterial culture for 10 minutes at 4500 rpm to obtain a pellet. We discarded the supernatant and resuspended the pellet in 500 μl of PBS. This solution was added to the sediment and shaken for five minutes to thoroughly mix the bacteria and the sediment. 1 ul aliquots of the solution were taken on D_0_ and every five days for one month and analyzed using a fluorescent microscope to check for E.coli survival and evidence of plasmid exchange. To further quantify the number of bacteria containing the plasmid, on D_0_ and D_30_ we plated out serial dilutions from 8 μg of soil in 1 mL of PBS to the 10^−4^ dilution on LB plates with kanamycin to quantify changes in CFU over time.

### Microscopy

To image transformed bacteria, mating experiments, and the soil survival experiment, we used a Zeiss AxioImager M1 microscope. To gain further insight into enzyme localization in *E.coli*, we performed structured illumination microscopy (SIM) using a Zeiss Elyra PS.1 Super Resolution microscope. We used cryotome sectioning and confocal microscopy using a Zeiss LSM710 to probe enzyme localization in EPS. For TEM, cell cultures were stained with 2% methanolic uranyl acetate and imaged using a Tecnai 10 TEM.

### Statistical analysis

We used one-way analysis of variance (ANOVA) to determine whether there were significant differences in microbial growth across the different treatments in the 96-well bioactivity assays. Differences among treatments were assessed by reference to the standard F tests. We used a limited number of t-tests to determine whether there were significant differences in growth between *P. putida* and the genetically engineered bacteria that grew the most in each assay. General statistics, ANOVA, and t-tests were conducted in R (v. 3.2.2, “Fire Safety”) using packages stats (v. 3.4, R core team) and psych (v. 1.6.4).

## Author contributions

KF created the vectors used in this study, performed the activity assays and microscopy, conducted the mating experiments, performed the statistical analysis of the data, and wrote the paper. ZZ conducted SPME GC/MS of cultures to detect dodecane and benzo(a)pyrene degradation. NT provided funding for this research.

## Acknowledgements

KF would like to thank Denise Schichnes of the Biological Imaging Facility at UC Berkeley for microscopy support, Danielle Jorgens from the Electron Microscopy Facility for help troubleshooting TEM, José Siles and Andrew Hendrickson for use of *P. citreus* and *M. oxydans* isolated from Shell Pond, and Michael Belcher for *P. putida* strain KT2440.

