## Supplementary Info for "Horizontal ‘gene drives’ harness indigenous bacteria for bioremediation"

### FIGURES AND TABLES

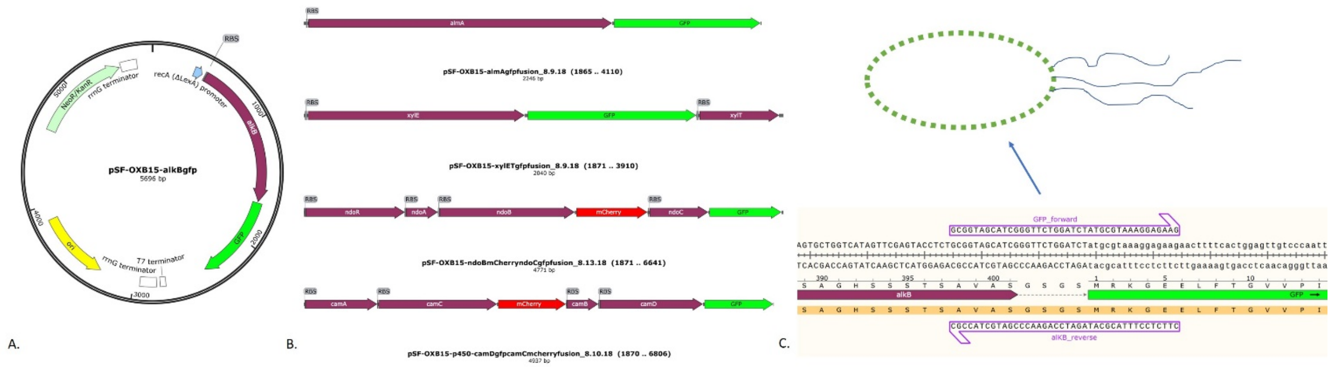


SI. Fig. 1 Overview of vector constructs. A. All constructs were made using the vector backbone pSF-OXB15, which contains the constitutive promoter recA and a kanamycin selection marker. B. 10 vectors were made in total; one vector for each enzyme containing the gene(s) of interest with a gfp marker included and one vector with gfp (and/or mcherry) fused to the mono or di-oxygenase (and if applicable, dehydrogenase) included in the vector. C. Overview of fusion protein system. Fluorophores were added to enzymes using five-seven glycines and serines.

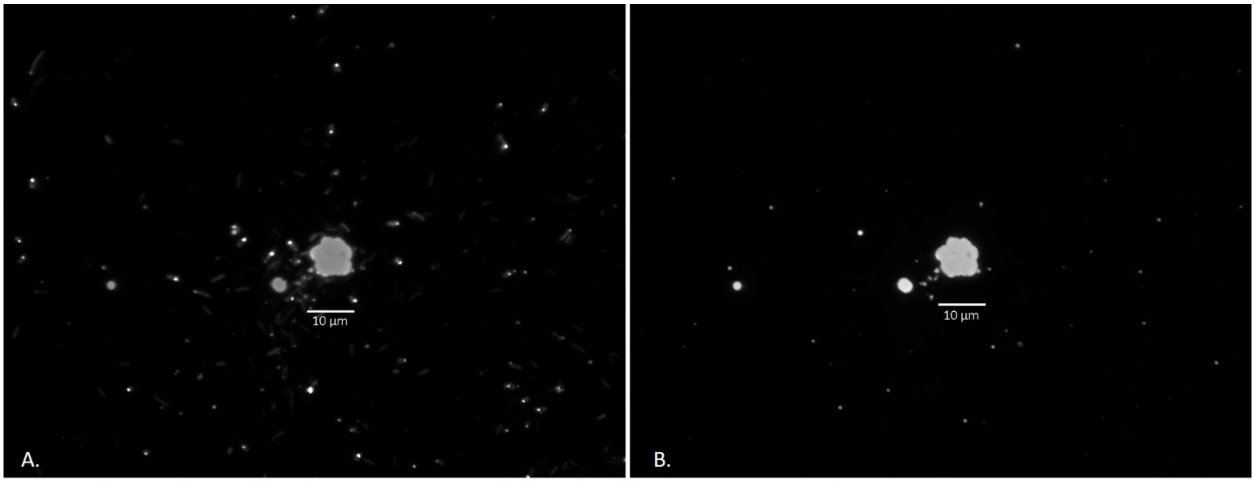


SI Fig. 2 Movement of E.coli expressing pSF-OXB15-xylefusion towards crude oil deposits. Bacteria are shown in the gfp channel in (A.) and crude oil stained with the fluorescent dye Nile Red is shown in the mcherry channel in (B.). E.coli were seen moving towards and attaching to oil droplets.

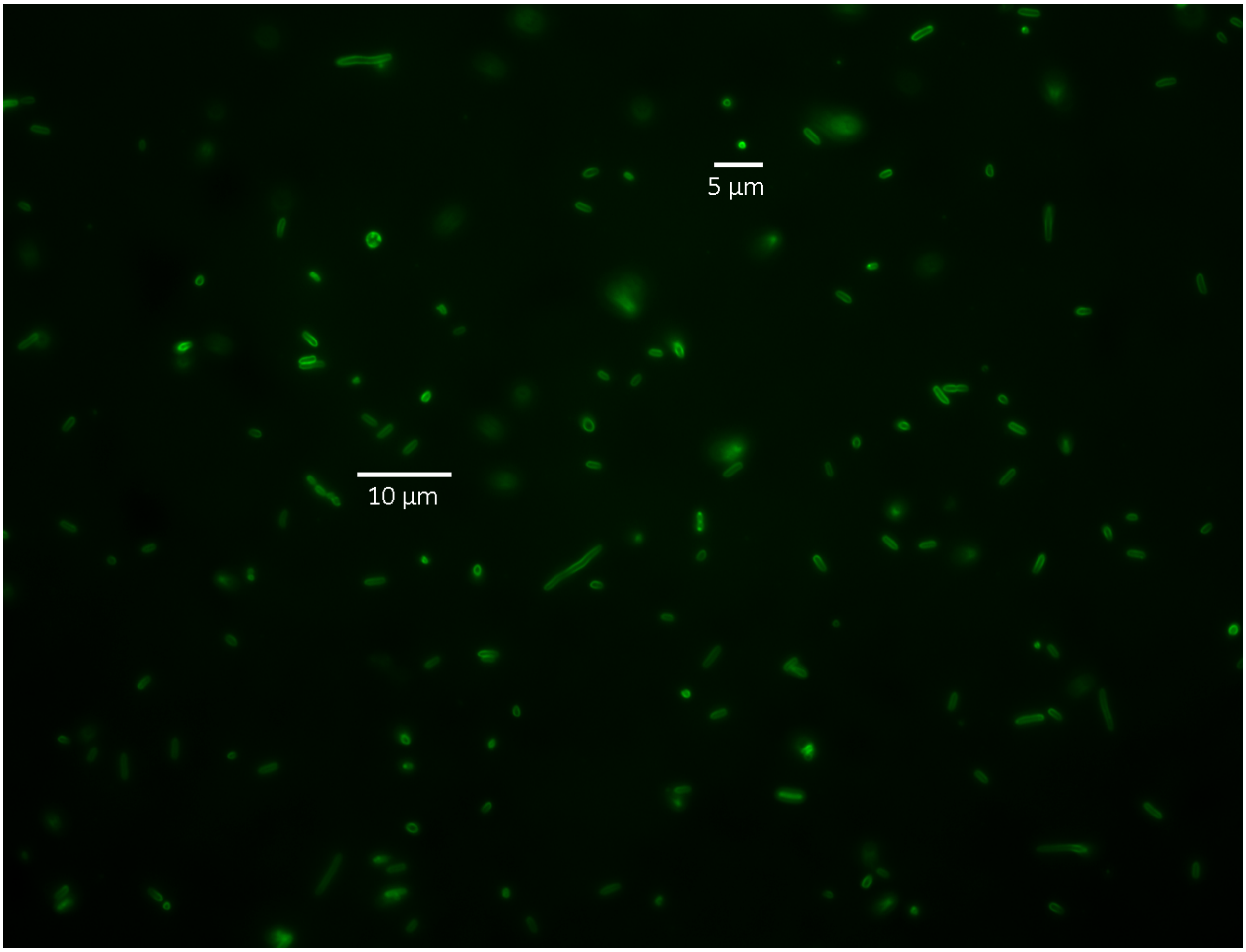


SI Fig. 3 Expression of alkB in E.coli. Here, alkB is fused to gfp, showing the localization of the enzyme to cell membranes and vesicles.

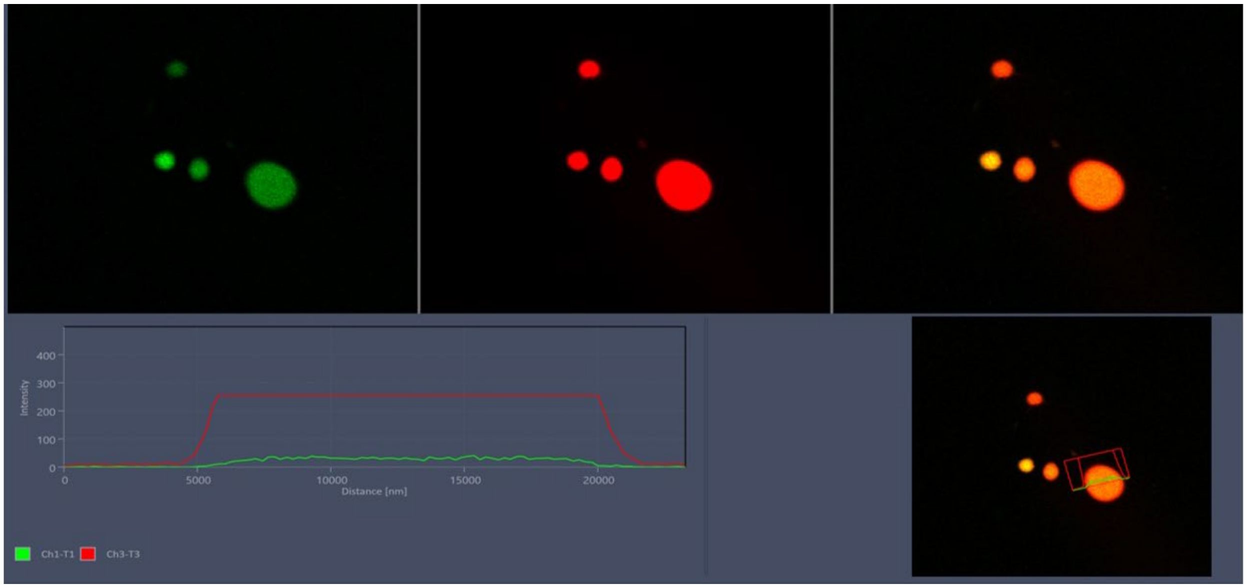


SI Fig. 4 Localization of extracellular enzymes to oil droplets. Here, oil droplets in cultures with E.coli expressing almA (fused to gfp) appear to be coated with the monooxygenase. The image shows a confocal image of a series of oil droplets showing gfp fluorescence (top left), mcherry fluorescence of the Nile Red-stained oil (top middle panel), and a composite of the two (far right top panel, yellow signal indicates overlap). The bottom panel shows a cross section of one of the oil droplets, confirming overlap of both fluorescent signals. In this case, the Nile Red fluorescence of the oil is far greater than the gfp signal coming from the enzyme almA.

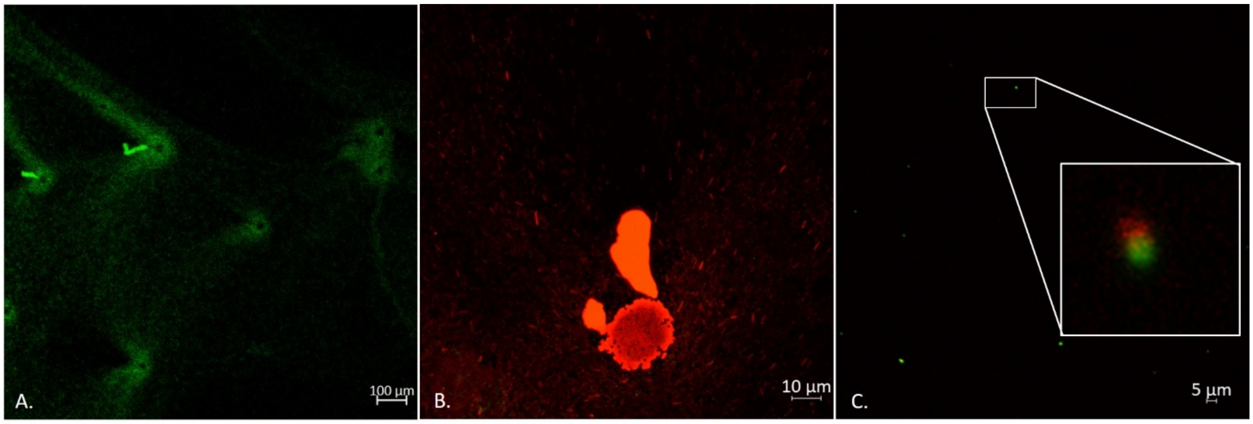


SI Fig. 5 Localization of p450camC and camD to E.coli EPS (from pSF-OXB15-p450camfusion). A. Confocal image (20x) of E.coli forming mounds and excreting EPS. B. Close-up confocal image (40x) of EPS excretion. C. Confocal image (100x) of cryotome sectioning of EPS. The box shows a magnified view of camC (red) and camD (green) localized to the same area. The two enzymes were frequently found together (roughly 60% of the time) but also found separately. In the box, each aggregate of enzyme is approximately 900 nm in diameter. Aggregates of protein varied considerably in size throughout the EPS but were always <1 um in diameter.

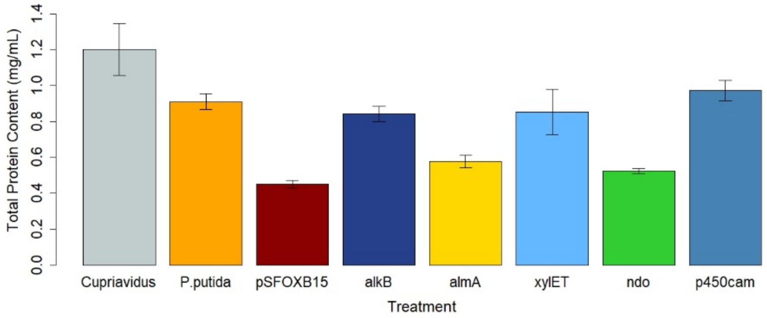


SI Fig. 6 Protein content from EPS produced by wild-type (*Cupriavidus* and *P. putida*), control (E.coli expressing the pSF-OXB15 backbone), and engineered bacteria. The engineered bacteria are designated by which enzyme they produced. Results are based on the non-fusion constructs.

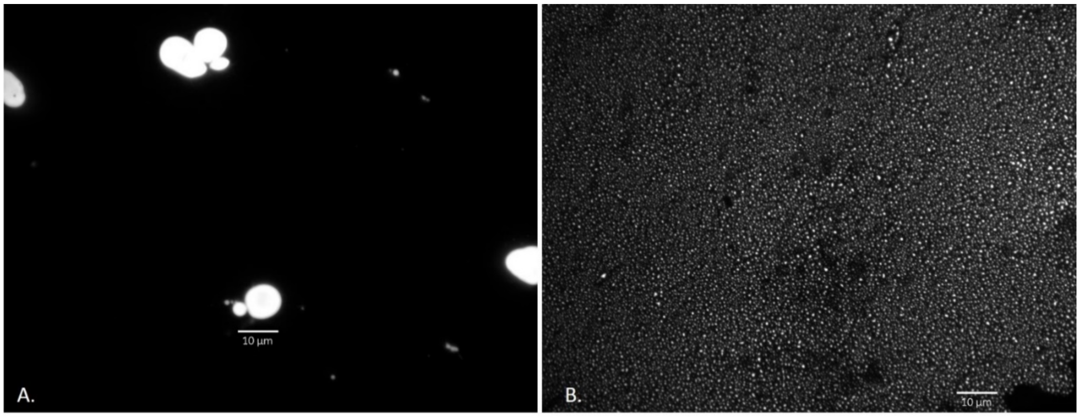


SI Fig. 7. Crude oil droplet formation appeared to vary among E.coli cultures containing different vectors. For example, the oil droplets in media with E.coli expressing alkB (A.) were larger than those from cultures expressing xylE (B.). At the moment, the relationship between enzyme expression and oil droplet formation is not clear.

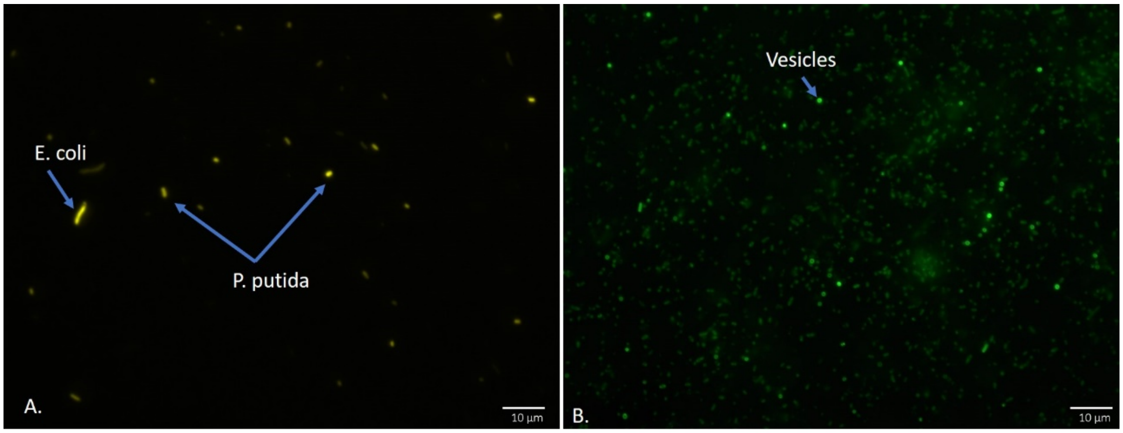


SI Fig. 8 Expression of pSF-OXB15-p450camfusion (A.) and alkB (B.) in *P.putida* after mating with *E.coli* harboring these genes.

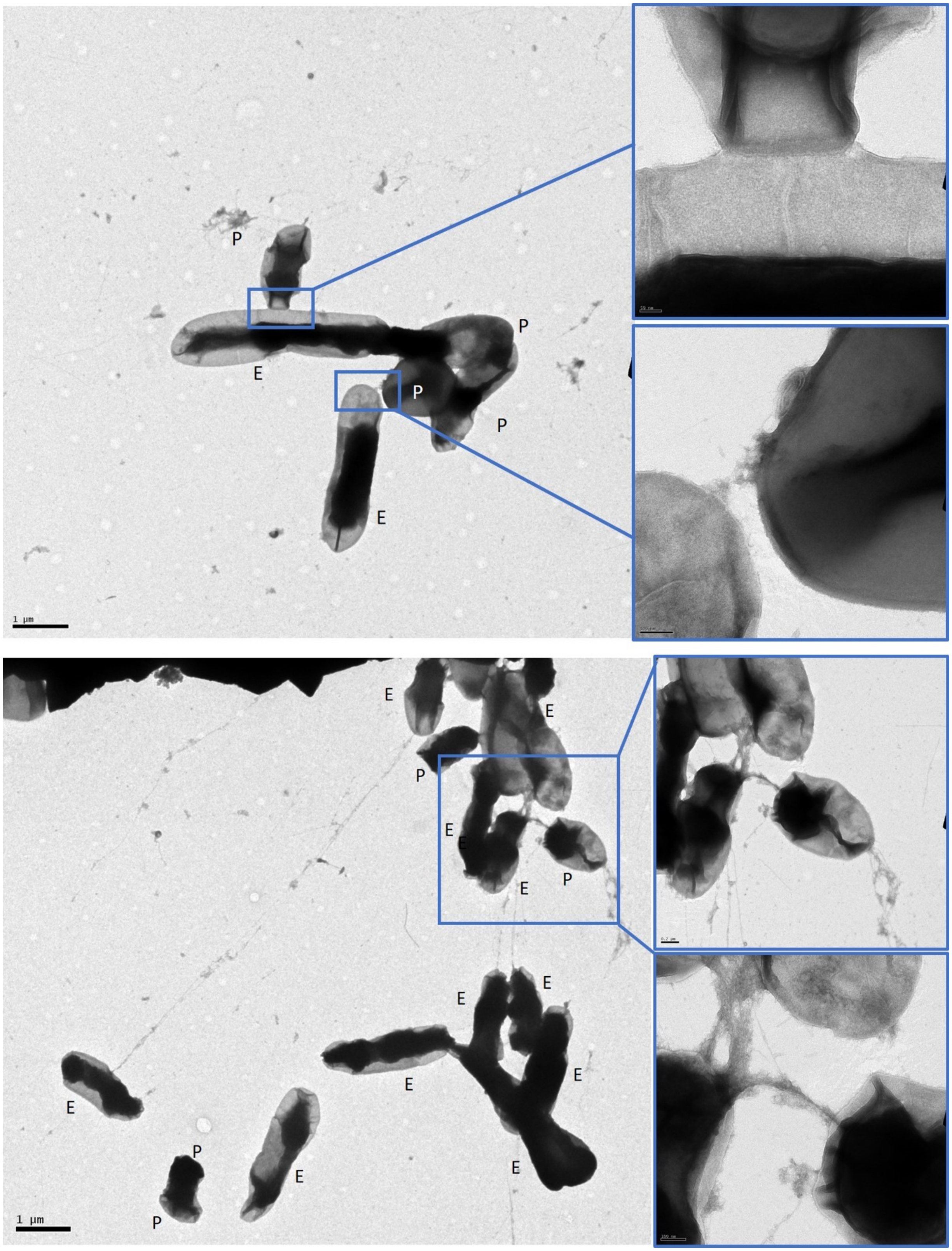


SI Fig. 9 TEM images of *E.coli* DH5α carrying pSF-OXB15-p450camfusion and *P. putida* connected by conjugative pilii, mating pair bridges, and nanotubes.

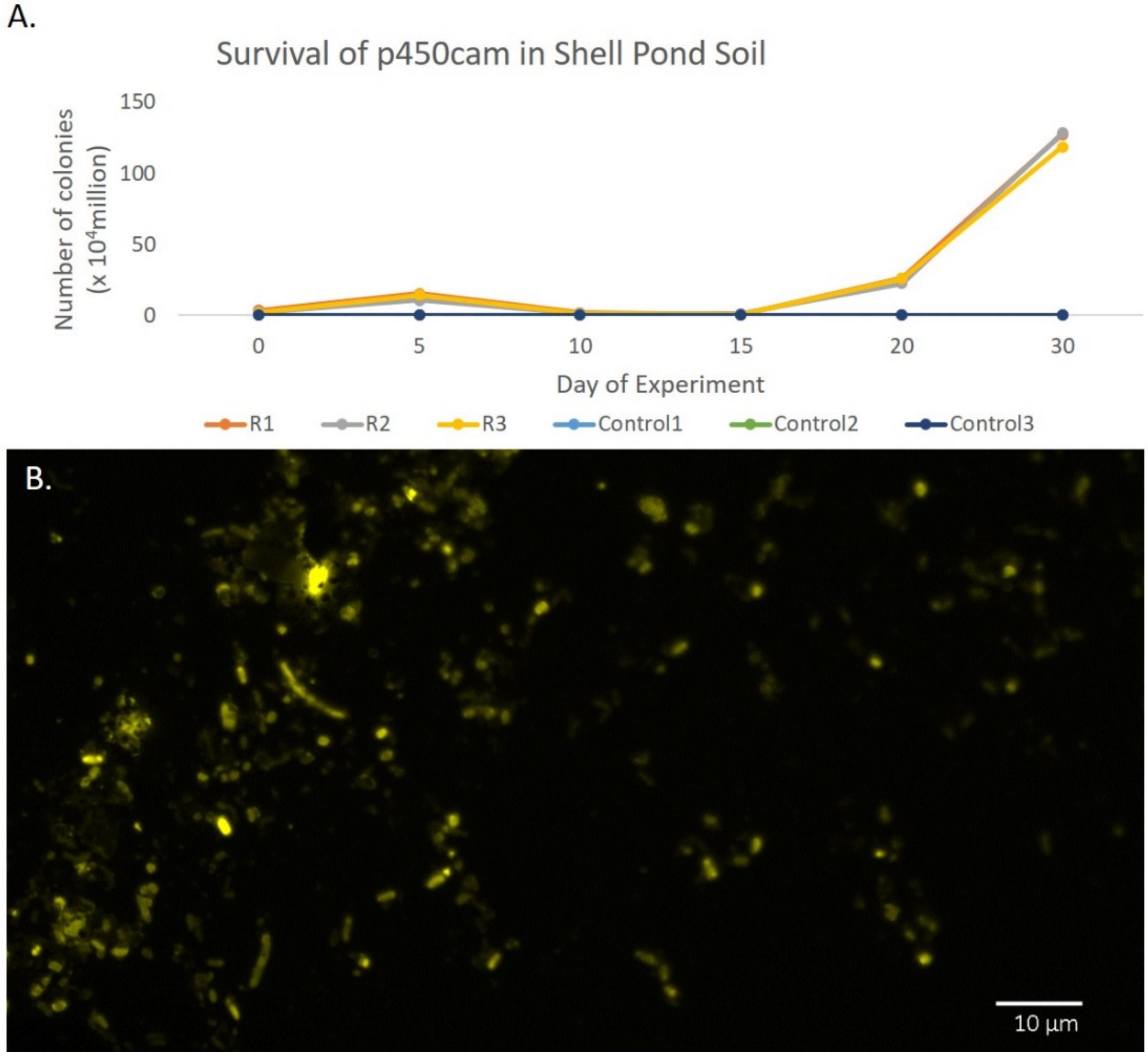


SI Fig. 10 Expression of vector pSF-OXB15-p450cammcherry in indigenous microbial communities from Shell Pond. A. Number of colonies on kanamycin plates from soil aliquots (8 µg) diluted from Shell Pond soil inoculated with *E. coli* DH5α harboring the plasmid pSF-OXB15-p450camfusion (R1-3) or no inoculation (control 1-3). The number of colonies declined from $D_{10}$-$D_{15}$ then continued to increase exponentially. B. Aliquot of Shell Pond soil showing diverse bacteria expressing the vector on $D_{21}$.

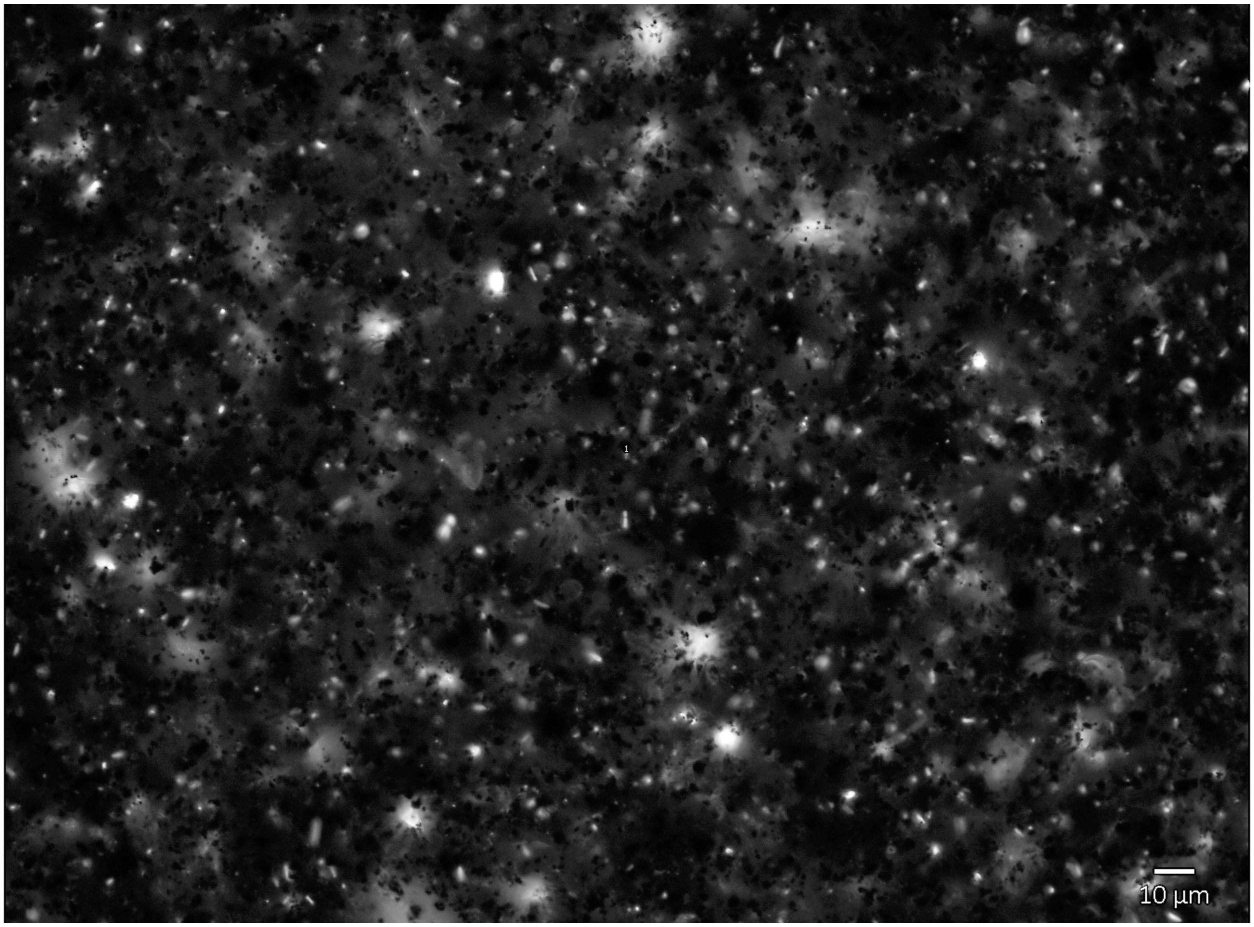


SI Fig. 11. Indigenous bacteria from Shell Pond carrying the plasmid pSF-OXB15-p450camfusion after 90 days. No re-inoculation was conducted during the course of the experiment.

**SI Table 1** Transformation rates for *P. putida*, *Cupriavidus* sp., *P. citreus*, and *M. oxydans* after 48h of exposure to E.coli expressing the vector pSF-OXB15-p450camfusion.

| Species | Total WT cells | Transformed WT cells | % transformed | Average Rate of Transformation |
|---|---|---|---|---|
| *P. putida* | 84 | 63 | 0.75 | **0.69** |
| *P. putida* | 57 | 41 | 0.72 | |
| *P. putida* | 52 | 32 | 0.62 | |
| *Cupriavidus* sp. | 106 | 43 | 0.41 | **0.67** |
| *Cupriavidus* sp. | 79 | 67 | 0.85 | |
| *Cupriavidus* sp. | 80 | 60 | 0.75 | |
| *P. citreus* | 51 | 42 | 0.82 | **0.84** |
| *P. citreus* | 19 | 17 | 0.89 | |
| *P. citreus* | 26 | 21 | 0.81 | |
| *M. oxydans* | 328 | 82 | 0.25 | **0.19** |
| *M. oxydans* | 425 | 67 | 0.16 | |

| | | | |
|---|---|---|---|
| *M. oxydans* | 338 | 58 | 0.17 |

**SI Table 2** Sequences of genes involved in petroleum hydrocarbon degradation used for the construction of the pSF-OXB15 series of plasmids used in this study.

| Gene | Sequence Source | Sequence |
|---|---|---|
| alkB | ENA AJ245436 | >atgcttgagaaacacagagttctggattccgctccagagtacgtagataaaaagaaatatctctggatactatcaactttgtggccggctactccgatgatcggaatctggcttgcaaatgaaactggttgggggatttttttatgggctggtattgctcgtatggtacggcgcacttccattgcttgatgcgatgtttggtgaggactttaataatccgcctgaagaagtggtgccgaaactagagaaggagcggtactatcgagttttgacatatctaacagttcctatgcattacgctgcattaattgtgtcagcatggtgggtcggaactcagccaatgtcttggcttgaaattggtgcgcttgccttgtcactgggtatcgtgaacggactagcgctcaatacaggacacgaactcggtcacaagaaggagacttttgatcgttggatggccaaaattgtgttggctgtcgtagggtacggtcacttctttattgagcataataagggtcatcaccgtgatgtcgctacaccgatggatcctgcaacatcccggatgggagaaagcatttataagttttcaatccgtgagatcccaggagcatttattcgtgcttgggggcttgaggaacaacgcctttcgcgccgtggccaaagcgtttggagtttcgataatgaaatcctccaaccaatgatcatcacagttattctttacgccgttctccttgccttgtttggacctaagatgctggtgttcctgccgattcaaatggctttcggttggtggcagctgaccagtgcgaactatattgaacattacggcttgctccgtcaaaaaatggaggacggtcgatatgagcatcaaaagccgcaccattcttggaatagtaatcacatcgtctctaatctagtgctgttccaccttcagcggcactcggatcaccacgcgcatccaacacgttcttatcagtcacttcgggattttcccggcctgccggctcttccgacgggttaccctggtgcatttttgatggcgatgattcctcagtggtttagatcagttatggatcccaaggtagtagattgggctggtggtgaccttaataagatccaaattgatgattcgatgcgagaaacctatttgaaaaaatttggcactagtagtgctggtcatagttcgagtacctctgcggtagcatcgtag |
| camA | NCBI AB771747 | >atgcacgtcaccctgcttgatacggcagcccgggttctggagcgggttaccgccccgccggtatcggccttttacgagcacctacaccgcgaagccggcgttgacatacgaaccggcacgcaggtgtgcgggttcgagatgtcgaccgaccaacagaaggttaccgccgtcctctgcgaggacggcacaaggctgccagcggatctggtaatcgccgggattggcctgataccaaactgcgagttggccagtgcggccggcctgcaggttgataacggcatcgtgatcaacgaacacatgcagacctctgatcccttgatcatggccgtcggcgactgtgcccgatttcacagtcagctctatgaccgctgggtgcgtatcgaatcggtgcccaatgccttggagcaggcacgaaagatcgccgccatcctctgtggcaaggtgccacgcgatgaggcggcgccctggttctggtccgatcagtatgagatcggattgaagatggtcggactgtccgaagggtacgaccggatcattgtccgcggctctttggcgcaacccgacttcagcgttttctacctgcagggagaccgggtattggcggtcgatacagtgaaccgtccagtggagttcaaccagtcaaaacaaataatcacggatcgtttgccggttgaaccaaacctactcggtgacgaaagcgtgccgttaaaggaaatcatcgccgccgccaaagctgaactgagtagtgcctga |
| camB | NCBI AB771747 | >atgtctaaagtagtgtatgtgtcacatgatggaacgcgtcgcgaactggatgtggcggatggcgtcagcctgatgcaggctgcagtctccaatggtatctacgatattgtcggtgattgtggcggcagcgccagctgtgccacctgccatgtctatgtgaacgaagcgttcacggacaaggtgcccgccgccaacgagcgggaaatcggcatgctggagtgcgtcacggccgaactgaagccgaacagcaggctctgctgccagatcatcatgacgcccgagctggatggcatcgtggtcgatgttcccgataggcaatggtaa |
| camC | NCBI AB771747 | >atgacgactgaaaccatacaaagcaacgccaatcttgcccctctgccaccccatgtgccagagcacctggtattcgacttcgacatgtacaatccgtcgaatctgtctgccggcgtgcaggaggcctgggcagttctgcaagaatcaaacgtaccggatctggtgtggactcgctgcaacggcggacactggatcgccactcgcggccaactgatccgtgaggcctatgaagattaccgccacttttccagcgagtgcccgttcatccctcgtgaagccggcgaagcctacgacttcattcccacctcgatggatccgcccgagcagcgccagtttcgtgcgctggccaaccaagtggttggcatgccggtggtggataagctggagaaccggatccaggagctggcctgctcgctgatcgagagcctgcgcccgcaaggacagtgcaacttcaccgaggactacgccgaacccttcccgatacgcatcttcatgctgctcgcaggtctaccggaagaagatatcccgcacttgaaatacctaacggatcagatgacccgtccggatggcagcatgaccttcgcagaggccaaggaggcgctctacgactatctgataccgatcatcgagcaacgcaggcagaagccgggaaccgacgctatcagcatcgttgccaacggccaggtcaatgggcgaccgatcaccagtgacgaagccaagaggatgtgtggcctgttactggtcggcggcctggatacggtggtcaatttcctcagcttcagcatggagttcctggccaaaagcccggagcatcgccaggagctgatcgagcgtcccgagcgtattccagccgcttgcgaggaactactccggcgcttctcgctggttgccgatggccgcatcctcacctccgattacgagtttcatggcgtgcaactgaagaaaggtgaccagatcctgctaccgcagatgctgtctggcctggatgagcgcgaaaacgcctgcccgatgcacgtcgacttcagtcgccaaaaggtttcacacaccacctttggccacggcagccatctgtgccttggccagcacctggcccgccgggaaatcatcgtcaccctcaaggaatggctgaccaggattcctgacttctccattgccccgggtgcccagattcagcacaagagcggcatcgtcagcggcgtgcaggcactccctctggtctgggatccggcgactaccaaagcggtataa |

| | | |
|---|---|---|
| camD | NCBI<br>AB771747 | >atgcaatacgcaagagcggctgtgatggttgagcaaaatcgggtcgagacctgggaggttccgattttcgatcctgcaccagg<br>tggggcactggtacgcgtggtgctaggcggcgtgtgcggaagcgacgtgcacatcgtatcaggggaggcaggcgccatgccctt<br>ccctatcattcttggccacgaaggcattggccgcatcgaaaagctcggcactggcgttaccacggattatgccggcgttccggtc<br>aaacagggcgacatggtctactgggcgcccattgcgctgtgccatcgctgccacagttgcaccgttctggatgaaacgccctgcg<br>acaactcgaccttttcgagcatgcgcaaaagcccaactggggcagttacgccgactttgcctgcctgcccaacggcatggcctt<br>ctaccggttgcccgatcatgcccagccagaagcgctggccgcgctcggttgcgccctaccgacagtactgcgcggctatgatcgc<br>tgtggcccggtcggcctggatgacaccgtcgtagtgcaaggcgcaggcccggtcggactcgcggccgtactggttgccgcagca<br>tccggcgccaaggacatcatcgccatagaccactccccgattcgcctggatatggcccgcagcctcggcgcgaccgaaaccatc<br>tccctggcagacaccacgccggaagagcgccagcggatcgtccaagaacgtttcggcaaacgtggcgccagcctggtcgtgga<br>agcggccggcgcgctaccagccttcccggaaggcgtgaatctgaccggcaaccacggccgctacgtcattctcgggctgtgggg<br>tgcaattggcacccagcccatctcgccccgggacttgaccatcaagaacatgagtattgccggagcaaccttcccgaagcccaa<br>gcattactaccaggccatgcagcttgcagctcgactgcaggatcgttatccgctggccgatctgatcacccagcgtttttccatcg<br>acgaggccagcaaggcacttgaactggtcaaggcaggagcactgatcaaacccgtgatcgaccccactctttag |
| almA | ENA<br>ABQ18224 | >atggaaaagcaagttgatgtattgattattggtgcaggtatctctgggattgggttagctgtacatctctctaaaaattgtccaca<br>acgcaaatttgaaatcttagagcgtcgtgaaagctttggtggaacttgggatttattccgctatcctggtattcgttcagattctgat<br>atgtcaacgtttggttttaacttcaagccatgggcaaaagacaaagtgttggctagcggtgctgagatcaaaggctatttaagtg<br>atgtgattagtgaaaatcagttaaaagacaaaattcactttggtcatcgtgtactttctgcgaactatgattctacaaagaaaaaa<br>tggttagttgagatcgaagataacaacaagaaaaaacaaacgtggtctgcaaacttcgtaatgggttgtacagggtactataac<br>tacgatcaaggttatgcacctaagttcccgaaacaagaagatttcaaaggtcagtttattcatccacagcactggccagaaaatt<br>tagattacacaggtaaaaaagttgtgatcatcggcagtggtgcaactgcaattacccttgtaccttctatggtcaaaggtggtgca<br>ggtcatgtgacgatgttacaacgttcgccaacgtatattgcgacgattccttctatcgactttatttatgaaaaaacacgtaaattt<br>atgtctgaagaaacggcttataaatttactcgtgcacgtaatattggtatgcaacgtggtatttatgcgctagcgcaaaagtatcc<br>aaaaactgtacgtcgtttattgttgaaaggcattgagttgcagttgaaaggcaaagtggatatgaaacactttactccaagttac<br>aatccttgggatcaacgtttatgtgttgtgccagatggtgatttattcaaagcattgcgtgaaggtcaggcaagtgttgagacaga<br>tcaaatcgagaaattcaccgcaaatggtattcaattgaaatctggtaaacatttagaggctgatattgtgatttcagcaactggtt<br>tagagattcagattttaggtggtgtccaaggttcaatcgatggtaaaccaatgaatacatctcaacatatgctttatcaaggcgtt<br>atggtgagcgatgtgcctaatatggcaatgatcattggttatatcaatgcgtcatggactttgaaggttgatattgctgctgattat<br>atttgccgtttgattaatcatatggataaaaatggttttgatgaggtgattgcacacgccgatccttcacagcgcgaaaatgacac<br>catcatgggtaaaatgtcatctggttatatcgcccgcgcagcggatgtgatgccaaaacaaggtaagcaagcaccttggaagat<br>cacaaataattaccttgcagatcgtaaggagttaaaagatgctaagtttaatgatggtgttttagaattccataagcgcggtgag<br>caaacagcaaaccgtaagccaaaattagtatcataa |
| xylE | ENA<br>AAA26052 | >atgaacaaaggtgtaatgcgaccgggccatgtgcagctgcgtgtactggacatgagcaaggccctggaacactacgtcgagtt<br>gctgggcctgatcgagatggaccgtgacgaccagggccgtgtctatctgaaggcttggaccgaagtggataagttttccctggtg<br>ctacgcgaggctgacgagccgggcatggattttatgggtttcaaggttgtggatgaggatgctctccggcaactggagcgggatc<br>tgatggcatatggctgtgccgttgagcagctacccgcaggtgaactgaacagttgtggccggcgcgtgcgcttccaggcccctc<br>cgggcatcacttcgagttgtatgcagacaaggaatatactggaaagtggggtttgaatgacgtcaatcccgaggcatggccgcg<br>cgatctgaaaggtatggcggctgtgcgtttcgaccacgccctcatgtatggcgacgaattgccggcgacctatgacctgttcacca<br>aggtgctcggtttctatctggccgaacaggtgctggacgaaaatggcacgcgcgtcgcccagtttctcagtctgtcgaccaaggc<br>ccacgacgtggccttcattcaccatccggaaaaaggccgcctccatcatgtgtccttccacctcgaaacctgggaagacttgcttc<br>gcgccgccgacctgatctccatgaccgacacatctatcgatatcggcccaacccgccacggcctcactcacggcaagaccatcta<br>cttcttcgacccgtccggtaaccgcaacgaagtgttctgcgggggagattacaactacccggaccacaaaccggtgacctggac<br>caccgaccagctgggcaaggcgatcttttaccacgaccgcattctcaacgaacgattcatgaccgtgctgacctga |
| xylT | ENA<br>AAA26051 | >atgaacagtgccggctacgaggtgttcgaagtgctaagcggccagtcattccgctgtgcggagggccagtcggtactgcgcgc<br>catggaagcccagggcaagcgctgcataccggtgggctgtcgcggtggcggttgcggcctttgtagagtgcgggtgctcagcgg<br>agcctaccggagcggacgcatgagccgcggtcacgtgccggccaaggccgccgccgaagcgctggccctggcctgtcaagtgtt<br>tccgcaaaccgacttgaccatcgagtactttcgccacgttggcggaaacaaacctgacaacatgaactatgaagaggtgacgtc<br>atga |
| ndoR | ENA<br>AAA25900 | >atggaacttctcatacagccaaacaatcgcctcattagctttagtcccggcgccaaccttctggaagtgcttcgcgaaaacggt<br>gtcgctatttcctacagttgtatgtctgggcgttgcggaacctgccgctgtcgggttacagatggcagtgtaattgattcgggggcg<br>ggaagcgggttaccaaacctcgtggacgagcattatgtgctcgcctgtcagtcagtacttactcacaattgcgcgatcgaaatcc<br>cagaaaccgacgaaatcgtcacccacccggcgagaatcatcaagggcactgtggtcgccgtcgagtcgcccactcacgatatcc<br>gtcgcctacgcgtacgcctcgctaagcccttcgagttctcacccggacagtacgcgacattgcagttcagtcctgagcatgcgcgt |

| | | |
|---|---|---|
| | | ccgtattcaatggcaggtctgccagatgaccaagaaatggagttccacatacgcaaggtgccgggtgggcgcgtaacggagtatgttttcgagcacgtccgcgaaggtacaagcatcaagttgagcgggccacttggtacggcttatttgcgtcagaaccacaccgggccgatgctctgtgtgggcggtgggaccggactagcaccggtgctgtcgattgttcgcggcgcgctgaagttgggtatgacaaacccccatcctcctttatttcggagtgcgcagtcagcaagacctctacgacgcagagcgattgcacaaactcgccgctgatcaccctcaactgaccgtacacacggtaatcgcaatgggcccgattaatgagagtcagcgagccggtctagttaccgatgtgatcgaaaaagacatcatttcgctggctgggtggagggcctacctgtgcggcgcaccagcgatggttgaagcgctttgcaccgttaccaagcatcttggaatatcacccgaacatatttatgccgatgccttctatcccggtggaatctga |
| ndoA | ENA AAB47590 | >atgacagtaaagtggattgaagcagtcgctctttctgacatccttgaaggtgacgtcctcggcgtgactgtcgagggcaaggagctggcgctgtatgaagttgaaggcgaaatctacgctaccgacaacctgtgcacgcatggttccgcccgcatgagtgatggttatctcgagggtagagaaatcgaatgccccttgcatcaaggtcggtttgacgtttgcacaggcaaagccctgtgcgcacccgtgacacagaacatcaaaacatatccagtcaagattgagaacctgcgcgtaatgattgatttgagctaa |
| ndoB | ENA AAB47591 | >atgaattacaataataaaatcttggtaagtgaatctggtctgagccaaaagcacctgattcatggcgatgaagaacttttccaacatgaactgaaaaccatttttgcgcggaactggctttttctcactcatgatagcctgattcctgcccccggcgactatgttaccgcaaaaatggggattgacgaggtcatcgtctcccggcagaacgacggttcgattcgtgcttttctgaacgtttgccggcatcgtggcaagacgctggtgagcgtggaagccggcaatgccaaaggttttgtttgcagctatcacggctggggcttcggctccaacggtgaactgcagagcgttccatttgaaaaagatctgtacggcgagtcgctcaataaaaaatgtctggggttgaaagaagtcgctcgcgtggagagcttccatggcttcatctacggttgcttcgaccaggaggcccctcctcttatggactatctgggtgacgctgcttggtacctggaacctatgttcaagcattccggcggtttagaactggtcggtcctccaggcaaggttgtgatcaaggccaactggaaggcacccgcggaaaactttgtgggagatgcataccacgtgggttggacgcacgcgtcttcgcttcgctcgggggagtctatcttctcgtcgctcgctggcaatgcggcgctaccacctgaaggcgcaggcttgcaaatgacctccaaatacggcagcggcatgggtgtgttgtgggacggatattcaggtgtgcatagcgcagacttggttccggaattgatggcattcggaggcgcaaagcaggaaaggctgaacaaagaaattggcgatgttcgcgctcggatttatcgcagccacctcaactgcaccgttttcccgaacaacagcatgctgacctgctcgggtgttttcaaagtatggaacccgatcgacgcaaacaccaccgaggtctggacctacgccattgtcgaaaaagacatgcctgaggatctcaagcgccgcttggccgactctgttcagcgaacgttcgggcctgctggcttctgggaaagcgacgacaatgacaatatggaaacagcttcgcaaaacggcaagaaatatcaatcaagagatagtgatctgctttcaaaccttggtttcggtgaggacgtatacggcgacgcggtctatccaggcgtcgtcggcaaatcggcgatcggcgagaccagttatcgtggtttctaccgggcttaccaggcacacgtcagcagctccaactgggctgagttcgagcatgcctctagtacttggcatactgaacttacgaagactactgatcgctaa |
| ndoC | ENA AAB47592 | >atgatgatcaatattcaagaagacaagctggtttccgcccacgacgccgaagagattcttcgtttcttcaattgccacgactctgctttgcaacaagaagccactacgctgctgacccaggaagcgcatttgttggacattcaggcttaccgtgcttggttagagcactgcgtggggtcagaggtgcaatatcaggtcatttcacgcgaactgcgcgcagcttcagagcgtcgttataagctcaatgaagccatgaacgtttacaacgaaaattttcagcaactgaaagttcgagttgagcatcaactggatccgcaaaactggggcaacagcccgaagctgcgctttactcgctttatcaccaacgtccaggccgcaatggacgtaaatgacaaagagctacttcacatccgctccaacgtcattctgcaccgggcacgacgtggcaatcaggtcgatgtcttctacgccgcccgggaagataaatggaaacgtggcgaaggtggagtacgaaaattggtccagcgattcgtcgattacccagagcgcatacttcagacgcacaatctgatggtctttctgtga |

**SI Table 3** Primers used in Gibson Assembly construction of plasmids used in this study.

| Primer number | Name | Sequence | Vector |
|---|---|---|---|
| 1 | alkB_Forward | TCCTACCATCCaggaggacagctATGCTTGAGAAACacagag | pSF_OXB15-alkBgfp |
| 2 | alkB_reverse | ctttacgcatagctgtcctcctctcCTACGATGCTACCG | pSF_OXB15-alkBgfp |
| 3 | gFP_Forward | TAGCATCGTAGGAGAGGAGGACAGCTATGCGTAAAGGAG | pSF_OXB15-alkBgfp |
| 4 | gfp_reverse | CAGTCAGTGCAGGAGGAGACAAttatttgtatagttcatcc | pSF_OXB15-alkBgfp |
| 5 | Vector_Forward | TTGTCTCCTCCTGCACTGACTGACTG | Vector backbone |
| 6 | Vector_Reverse | agctgtcctcctGGATGGTAGGATCGACAAAG | Vector backbone |
| 7 | alkB_Forward | TCCTACCATCCaggaggacagctATGCTTGAGAAACacagag | pSF-OXB15-alkBgfpfusion |
| 8 | alKB_reverse | CTTCTCCTTTACGCATAGATCCAGAACCCGATGCTACCGC | pSF-OXB15-alkBgfpfusion |

| | | | |
|---|---|---|---|
| **9** | gfp_forward | GCGGTAGCATCGGGTTCTGGATCTATGCGTAAAGGAGAAG | pSF-OXB15-alkBgfpfusion |
| **10** | gfp_reverse | CAGTCAGTGCAGGAGGAGACAAttatttgtatagttcatcc | pSF-OXB15-alkBgfpfusion |
| **11** | Vector_Forward | TTGTCTCCTCCTGCACTGACTGACTG | Vector backbone |
| **12** | Vector_Reverse | agctgtcctcctGGATGGTAGGATCGACAAAG | Vector backbone |
| **23** | pSF-OXB15_fwd | TTGTCTCCTCCTGCACTGACTGACTG | Vector backbone |
| **24** | pSF-OXB15_rev | agctgtcctcctGGATGGTAGGATCGACAAAG | Vector backbone |
| **25** | camA_fwd | atcctaccatccaggaggacagctATGCACGTCAC | pSF-OXB15-p450cam |
| **26** | camA_rev | cagtcgtcatagctgtcctcctctcTCAGGCACTACTCAGTTC | pSF-OXB15-p450cam |
| **27** | camB_fwd | agcggtataacacaggaggacagctATGTCTAAAGTAGTGTATG | pSF-OXB15-p450cam |
| **28** | camB_reverse | TTGCATAGCTGTCCTCCTGTAAGATTACCATTGCCTATC | pSF-OXB15-p450cam |
| **29** | camC_fwd | tagtgcctgagagaggaggacagctATGACGACTGAAACC | pSF-OXB15-p450cam |
| **30** | camC_reverse | ctactttagacatagctgtcctcctgtgttataccgctttgg | pSF-OXB15-p450cam |
| **31** | camD_fwd | aatggtaatcttacaggaggacagctATGCAATACGCAAGAG | pSF-OXB15-p450cam |
| **32** | camD_reverse | ctttacgcatagctgtcctcctctagtcctaaagagtggggtcgatc | pSF-OXB15-p450cam |
| **33** | gfp_fwd | CACTCTTTAGGACTAGAGGAGGACAGCTATGCGTAAAGGAGAAGAAC | pSF-OXB15-p450cam |
| **34** | gfp_rev | CAGTCAGTGCAGGAGGAGACAAttatttgtatagttcatcc | pSF-OXB15-p450cam |
| **35** | pSF-OXB15_fwd | TTGTCTCCTCCTGCACTGACTGACTG | Vector backbone |
| **36** | Vector_Reverse | agctgtcctcctGGATGGTAGGATCGACAAAG | Vector backbone |
| **37** | camA_fwd | atcctaccatccaggaggacagctATGCACGTCAC | pSF-OXB15-450cam_fusion |
| **38** | camA_rev | cagtcgtcatagctgtcctcctctcTCAGGCACTACTCAGTTC | pSF-OXB15-450cam_fusion |
| **39** | camB_forward | tgtacaagTAAcacaggaggacagctatgtctaaagtag | pSF-OXB15-450cam_fusion |
| **40** | camB_reverse | TTGCATAGCTGTCCTCCTGTAAGATTACCATTGCCTATC | pSF-OXB15-450cam_fusion |
| **41** | camC_fwd | tagtgcctgagagaggaggacagctATGACGACTGAAACC | pSF-OXB15-450cam_fusion |
| **42** | camC_reverse | TGCTCACCATGCCAGATCCTGAACCACCTACCGCTTTGG | pSF-OXB15-450cam_fusion |
| **43** | camD_fwd | aatggtaatcttacaggaggacagctATGCAATACGCAAGAG | pSF-OXB15-450cam_fusion |
| **44** | camD_reverse | TTACGCATAGAACCCGAGCCAGATCCGCTAAGAGTGGGGTCG | pSF-OXB15-450cam_fusion |
| **45** | gfp_forward | TCGACCCCACTCTTAGCggaTCTggcTCGGGTTCTATGCGTAAAGGAG | pSF-OXB15-450cam_fusion |
| **46** | gfp_rev | CAGTCAGTGCAGGAGGAGACAAttatttgtatagttcatcc | pSF-OXB15-450cam_fusion |
| **47** | mCherry_forward | accaaagcggtaggtggttcaggatctggcatggtgagcaag | pSF-OXB15-450cam_fusion |
| **48** | mCherry_reverse | CTTTAGACATAGCTGTCCTCCTGTGTTACTTGTACAGC | pSF-OXB15-450cam_fusion |
| **49** | pSF-OXB15_fwd | TTGTCTCCTCCTGCACTGACTGACTG | Vector backbone |
| **50** | Vector_Reverse | agctgtcctcctGGATGGTAGGATCGACAAAG | Vector backbone |
| **51** | almA_forward | gatcctaccatccaggaggacagctATGGAGAAACAGGTGGACG | pSF-OXB15_almgfpOPT |
| **52** | almA_rev | tcctttacgcatagcagtcctcctcgggagTCAGCTAACCAACTTCGGTTTG | pSF-OXB15_almgfpOPT |
| **53** | gfp-reverse | CAGTCAGTGCAGGAGGAGACAAttatttgtatagttcatcc | pSF-OXB15_almgfpOPT |

| | | | |
|---|---|---|---|
| **54** | gfp_fwd | ttggttagctgactcccgaggaggactgctATGCGTAAAGGAGAAGAAC | pSF-OXB15_almgfpOPT |
| **55** | pSF-OXB15_fwd | TTGTCTCCTCCTGCACTGACTGACTG | Vector backbone |
| **56** | Vector_Reverse | agctgtcctcctGGATGGTAGGATCGACAAAG | Vector backbone |
| **57** | almA_rev | tctcctttacgcatAGATCCAGAACCGCTAACCAACTTCGG | p-SF-OXB15-AlmAgfpfusionOPT |
| **58** | almA_fwd | gatcctaccatccaggaggacagctATGGAGAAACAGGTGGACG | p-SF-OXB15-AlmAgfpfusionOPT |
| **59** | gfp-Forward | CGAAGTTGGTTAGCGGTTCTGGATCTatgcgtaaaggagaag | p-SF-OXB15-AlmAgfpfusionOPT |
| **60** | gfp-reverse | CAGTCAGTGCAGGAGGAGACAAttatttgtatagttcatcc | p-SF-OXB15-AlmAgfpfusionOPT |
| **61** | pSF-OXB15_fwd | TTGTCTCCTCCTGCACTGACTGACTG | Vector backbone |
| **62** | Vector_Reverse | agctgtcctcctGGATGGTAGGATCGACAAAG | Vector backbone |
| **63** | gfp-reverse | CAGTCAGTGCAGGAGGAGACAAttatttgtatagttcatcc | pSF-OXB15-xylETgfp |
| **64** | gfp_forward | GACGAGCTGACACAGGAGGTAGTATATGCGTAAAG | pSF-OXB15-xylETgfp |
| **65** | pSF-OXB15_fwd | TTGTCTCCTCCTGCACTGACTGACTG | pSF-OXB15-xylETgfp |
| **66** | Vector_Reverse | agctgtcctcctGGATGGTAGGATCGACAAAG | pSF-OXB15-xylETgfp |
| **67** | xylE_forward | ATCCTACCATCCAGGAGGACAGCTATGAACAAAGGTG | pSF-OXB15-xylETgfp |
| **68** | xylE_rev | cactgttcatagctgtcctcgtctcTCAGGTCAGCACG | pSF-OXB15-xylETgfp |
| **69** | xylT_fwd | GCTGACCTGAGAGACGAGGACAGCTATGAACAGTGC | pSF-OXB15-xylETgfp |
| **70** | xylT_reverse | ctttacgcatatactacctcctgtgtcagctcgtcacctc | pSF-OXB15-xylETgfp |
| **71** | gfp-forward | cgtgctgaccagtGGTTCTGGATCTatgcgtaaaggag | pSF-OXB15-XylEGFPfusion |
| **72** | gfp_reverse | CACTGTTCATAGCTGTCCTCGTCTCTTATTTGTATAGTTC | pSF-OXB15-XylEGFPfusion |
| **73** | pSF-OXB15_fwd | TTGTCTCCTCCTGCACTGACTGACTG | pSF-OXB15-XylEGFPfusion |
| **74** | Vector_Reverse | agctgtcctcctGGATGGTAGGATCGACAAAG | pSF-OXB15-XylEGFPfusion |
| **75** | xylE_forward | ATCCTACCATCCAGGAGGACAGCTATGAACAAAGGTG | pSF-OXB15-XylEGFPfusion |
| **76** | xylE_reverse | ctttacgcatAGATCCAGAACCactggtcagcacggtc | pSF-OXB15-XylEGFPfusion |
| **77** | xylT_forward | ctatacaaataagagacgaggacagctatgaacagtgctg | pSF-OXB15-XylEGFPfusion |
| **78** | xylT_reverse | GCAGGAGGAGACAATCAGCTCGTCACC | pSF-OXB15-XylEGFPfusion |
| **79** | gfp_forward | TCTTTCTGTGACAGCATAGGAGGACAAGTATGCGTAAAG | pSF-OXB15-ndo |
| **80** | gfp_rev | CAGTCAGTGCAGGAGGAGACAAttatttgtatagttcatcc | pSF-OXB15-ndo |
| **81** | ndoA_forward | tggaatctgagagaggaggacagctatgacagtaaagtggattg | pSF-OXB15-ndo |
| **82** | ndoA_reverse | tattgtaattcatagctgtcctccttgtcggttagctcaaatcaatcattac | pSF-OXB15-ndo |
| **83** | ndoB_forward | GATTTGAGCTAACCGACAAGGAGGACAGCTATGAATTACAATAATAAAATC | pSF-OXB15-ndo |
| **84** | ndoB_reverse | ttatcatcatagctgtcctcctctgtctttagcgatcagtagtcttc | pSF-OXB15-ndo |
| **85** | ndoC | ctgatcgctaaagacagaggaggacagctatgatgataaatattc | pSF-OXB15-ndo |
| **86** | ndoC_reverse | tttacgcatacttgtcctcctatgctgtcacagaaagaccatc | pSF-OXB15-ndo |
| **87** | ndoR_fwd | gatcctaccatccaggaggacagctATGGAACTTCTCATAC | pSF-OXB15-ndo |
| **88** | ndoR_Reverse | ctttactgtcatagctgtcctcctctctcagattccaccgggatag | pSF-OXB15-ndo |
| **89** | pSF-OXB15_forward | TTGTCTCCTCCTGCACTGACTGACTG | Vector backbone |

| | | | |
|---|---|---|---|
| **90** | Vector_Reverse | agctgtcctcctGGATGGTAGGATCGACAAAG | Vector backbone |
| **91** | gfp_forward | ggtctttctgagtggctctggatctatgcgtaaag | pSF-OXB15-ndoBCfusion |
| **92** | gfp_rev | CAGTCAGTGCAGGAGGAGACAAttatttgtatagttcatcc | pSF-OXB15-ndoBCfusion |
| **93** | mCherry_forward | ctactgatcgcagtgggagttctggaggcatggtgagcaag | pSF-OXB15-ndoBCfusion |
| **94** | mcherry_reverse | tttatcatcatacctgtcctcctgcctctTTActtgtacagctcgtc | pSF-OXB15-ndoBCfusion |
| **95** | ndoA_forward | tggaatctgagagaggaggacagctatgacagtaaagtggattg | pSF-OXB15-ndoBCfusion |
| **96** | ndoA_reverse | tattgtaattcatagctgtcctccttgtcggttagctcaaatcaatcattac | pSF-OXB15-ndoBCfusion |
| **97** | ndoB_forward | GATTTGAGCTAACCGACAAGGAGGACAGCTATGAATTACAATAATAAAATC | pSF-OXB15-ndoBCfusion |
| **98** | ndoB_reverse | TGCTCACCATGCCTCCAGAACTCCCACTGCGATCAG | pSF-OXB15-ndoBCfusion |
| **99** | ndoC_forward | gctgtacaagtaaagaggcaggaggacaggtatgatgataaatattc | pSF-OXB15-ndoBCfusion |
| **100** | ndoC_reverse | tttacgcatagatccagagccactcagaaagaccatcag | pSF-OXB15-ndoBCfusion |
| **101** | ndoR_fwd | gatcctaccatccaggaggacagctATGGAACTTCTCATAC | pSF-OXB15-ndoBCfusion |
| **102** | ndoR_Reverse | ctttactgtcatagctgtcctcctctctcagattccaccgggatag | pSF-OXB15-ndoBCfusion |
| **103** | pSF-OXB15_forward | TTGTCTCCTCCTGCACTGACTGACTG | Vector backbone |
| **104** | Vector_Reverse | agctgtcctcctGGATGGTAGGATCGACAAAG | Vector backbone |
